# Quantitative understanding of molecular competition as a hidden layer of gene regulatory network

**DOI:** 10.1101/258129

**Authors:** Ye Yuan, Lei Wei, Tao Hu, Shuailin Li, Tianrun Cheng, Jinzhi Lei, Zhen Xie, Michael Q. Zhang, Xiaowo Wang

## Abstract

Molecular competition is ubiquitous, essential and multifunctional throughout diverse biological processes. Competition brings about trade-offs of shared limited resources among the cellular components, and it thus introduce a hidden layer of regulatory mechanism by connecting components even without direct physical interactions. By abstracting the analogous competition mechanism behind diverse molecular systems, we built a unified coarse-grained competition motif model to systematically compare experimental evidences in these processes and analyzed general properties shared behind them. We could predict in what molecular environments competition would reveal threshold behavior or display a negative linear dependence. We quantified how competition can shape regulator-target dose-response curve, modulate dynamic response speed, control target expression noise, and introduce correlated fluctuations between targets. This work uncovered the complexity and generality of molecular competition effect, which might act as a hidden regulatory mechanism with multiple functions throughout biological networks in both natural and synthetic systems.

## Introduction

Competition for limited resources matters at all scales of biology. Competition among different species can alter population distributions and ecological niches (Connell, 1983; Hardin, 1960; Schoener, 1983). Competition among individuals of the same species may slow down the growth rates of all competitors, driving natural selection and evolution (Bolnick, 2004; Svanback & Bolnick, 2007; Zwietering et al., 1990). Competition among adjacent cells in an organism can regulate their growth and viability, and enhance the dominance of cells with better fitness (Chang et al., 2015; Johnston, 2009; Khare & Shaulsky, 2006; Laird, 1964). In a microscopic scale, biological molecules within cells also face competition. Competition brings about trade-offs of shared limited resources among the cellular components (Hui et al., 2015; Scott et al., 2010; Weisse et al., 2015), and it thus introduces a hidden layer of regulatory mechanism by connecting components even without direct physical interactions. Miscellaneous phenomena caused by molecular competition have been reported in a variety of biological processes in diverse organisms. For example, DNA binding sites on plasmids can compete for transcription factor (TF) LacI to dictate its target gene expression in *E. coli* (Brewster et al., 2014). Noncoding RNAs transcribed from enhancer or promoter region can competitively bind to TF Yin-Yang 1 to trap the TF locally thus maintain gene expression stability in mouse embryonic stem cells (Sigova et al., 2015). mRNA, long-noncoding RNA and circular RNA molecules can competitively bind to microRNAs (miRNAs) to regulate various processes, such as cell growth (Zheng et al., 2016), cell differentiation (Cesana et al., 2011) and tumor suppression (Sumazin et al., 2011). Competition between RNA binding proteins PGL-3 and MEX-5 for mRNA drives polar positioning of phase-separated liquid compartments in *C. elegans* embryos (Saha et al., 2016). Furthermore, competition effects are especially important in synthetic gene circuits. Every synthetic gene inevitably competes for common resources with each other in circuits and with endogenous biological processes, introducing unexpected circuit failures or host metabolic burdens (Cardinale & Arkin, 2012; Qian et al., 2017; Wu et al., 2016). In addition, when one genetic element drives two or more downstream elements, competition will modulate the dynamics of signal transduction (Jayanthi et al., 2013; Jiang et al., 2011). As a result, characteristics of each single component are insufficient for the accurate prediction of the whole circuit behavior, posing a serious obstacle in synthetic circuit design and application.

Several mathematical frameworks and synthetic gene experiments have been built to quantitatively understand the diverse biological phenomena caused by competition. For example, a thermodynamic model was used to explain the TF titration effect in *E. coli* (Brewster et al., 2014). Kinetic model has been adopted to analyze competing endogenous RNA (ceRNA) regulation (Ala et al., 2013), and we further quantified the ceRNA effect through synthetic gene circuits in human cell line (Yuan et al., 2015). A minimal model based on delay differential equations was established to describe ribosome allocation between endogenous and synthetic genes in *E. coli* (Gorochowski et al., 2016). Queueing theory was introduced to describe the protein degradation process in *E. coli,* where target proteins as queues compete for degradation machine ClpXP as server (Cookson et al., 2011; Mather et al., 2010). However, common properties and underlying competition mechanisms in essence behind these diverse phenomena have not been systematically analyzed yet.

Here we propose that regulations by competition are ubiquitous, essential and multifunctional through diverse biological regulatory processes. By abstracting the analogous competition motif shared by diverse molecular systems, we built a unified coarse-grained kinetic model to systematically integrate experimental evidences in diverse biological processes and analyze the common properties shared among them. We organized these properties from steady-state behavior to dynamic responses, to quantify how competition could introduce constraints and indirect regulations among the targets and how the existence of competitors might influence regulator-target response characteristics. This work demonstrated the complexity and generality of the molecular competition effect, which is a ubiquitous hidden regulatory mechanism with diverse functions throughout different biological processes in both natural and synthetic life systems.

## Results

### A unified coarse-gained competition motif model

A number of phenomena caused by molecular competition have been reported in diverse biological systems recently (Brewster et al., 2014; Saha et al., 2016; Sigova et al., 2015; Zheng et al., 2016). Do they share any common properties? Could they be described by a unified model? We summed up several representative competition scenarios following the life cycle of gene expression (Figure 1), including competitions for transcription factors by DNA binding sites (Figure 1B), competitions for miRNAs and ribosomes by RNA molecules (Figure 1C and 1D), and competitions for degradation enzymes by target proteins (Figure 1E). Inspired by previous models studying ceRNA effect (Ala et al., 2013; Yuan et al., 2015), we proposed a generalized competition motif model, in which two target molecule species (target#1 and #2, and *T*_2_) competitively bind with a shared regulatory molecule species (regulator, R) (Figure 1A), to describe the similar competition topology these cases share. In this model, each molecule species is produced and degraded with certain rates, and the regulator is dynamically bound to targets following biochemical mass-action laws to form complexes (Figure 1F, SI Material and Methods). Loss rates of regulator (α) and its competing targets (*β*) were introduced to describe reactions from pure stoichiometric (α ~ 1, *β* ~ 1) to pure catalytic (α ~ 1, *β* ~ 0 where enzymes act as competitors, or *α* ~ 0, *β* ~ 1 when substrates act as competitors) (Ala et al., 2013). In different biochemical scenarios, experimentally measured signals may reflect different component levels of the competition motif. For example, the activity of targets could be mainly reflected by the abundance of complexes (*T^C^*) when the regulator is an activator, or by the abundance of the free targets (*T^F^*) when the regulator is a repressor.

**Figure 1.**
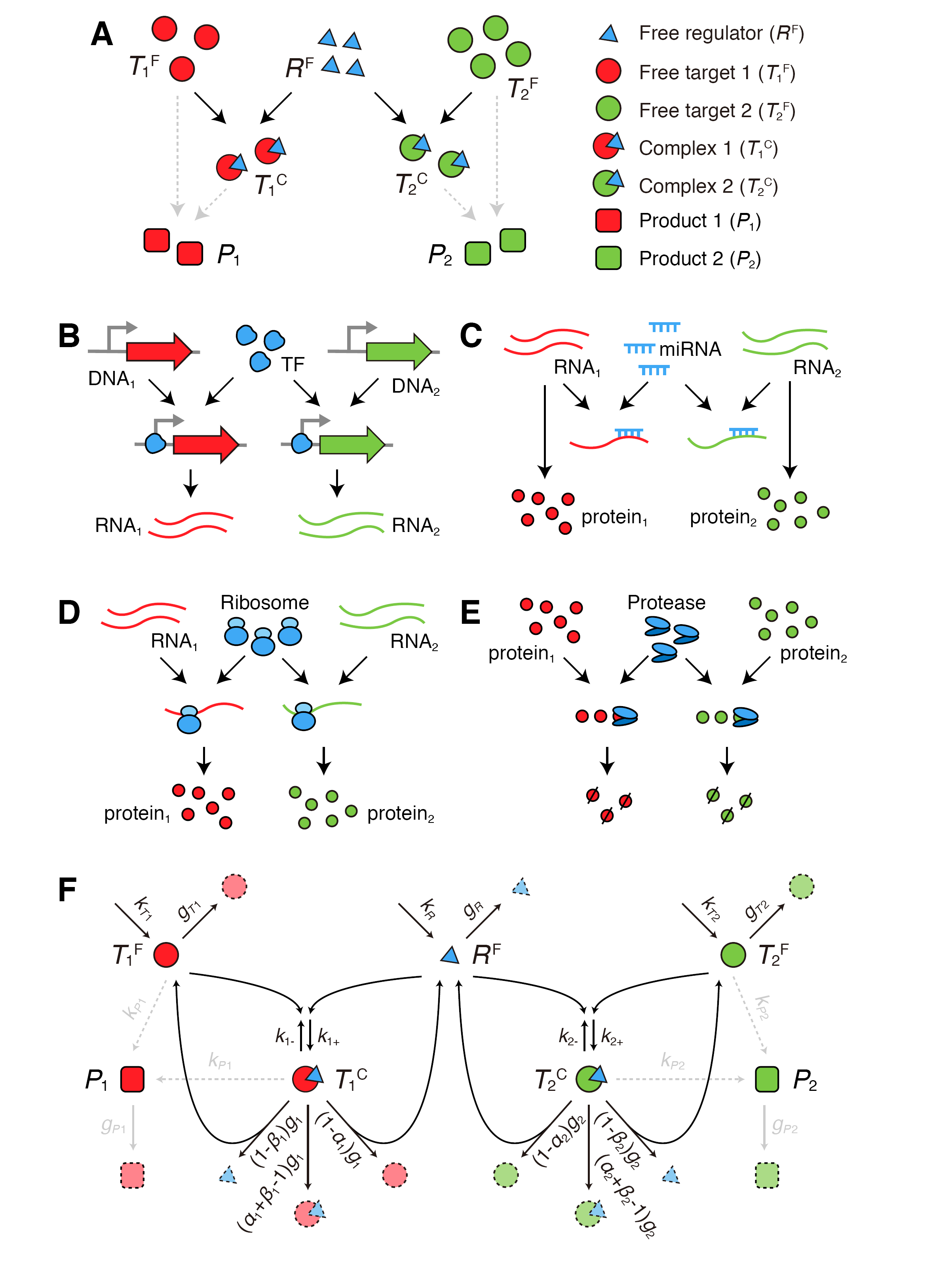
The coarse-gained competition motif model.(**A**) Basic structure of the competition motif. Downstream products can be produced from either free targets or complexes.(**B-E**) Competition motifs abstracted from diverse competition scenarios: (**B**) DNA binding sites competing for TFs; (**C**) RNA molecules competing for miRNAs; (**D**) mRNA molecules competing for ribosomes; (**E**) proteins competing for proteases.(**F**) Unified kinetic model of the competition motif.

This unified model can describe competitions in various biological processes (Figure S1A-D). Despite of different parameter settings, all these cases share the core competition motif structure, suggesting that they may share common characteristics. In the following sections, we used this model to analyze, in the scenario of either steady-state behavior or dynamic response, how the competition introduces indirect regulations between targets and how the existence of the competitors influences the property of regulator-target response.

### Relative abundance determines the regulatory patterns between competitors

Competition can cause crosstalk between targets. By quantifying the competition effect of one target upon the abundance of another target, recent studies have reported two apparently different steady-state behaviors named “threshold behavior” of ceRNA regulation in mammalian cells (Ala et al., 2013) and “negative linear dependence” behavior of synthetic gene expression in bacteria (Carbonell-Ballestero et al., 2016; Gyorgy et al., 2015). How could competition generate such two vastly different phenomena?

The model predicted that the relative abundance between regulator and target determines the diverse behaviors. Figure 2A and S2A illustrates how molecular abundance changes along with the gradual increment of *T_2_’s* production rate. The system went through three regimes: *“R* abundant”, “R near-equimolar” and *“R* scarce”, which are mainly determined by the production rate and loss rate of each component (SI Material and Methods). In the “R abundant” regime, free *T*_1_ level (*T_1_^F^*) is not sensitive to the increment of free *T*_2_ level (*T*_2_^F^), but when the system enters the “*R* near-equimolar” regime, *T*_1_^F^ becomes more sensitive to *T*_2_^F^ changes, thus generates the threshold behavior (Figure 2B and S2B). In contrast, *T*_1_ complex level (*T*_1_^C^) is substantially unchanged with respect to *T*2 complex level (*T*2^C^) except in the *“R* scarce” regime, where *T*_1_^C^ displays a negative linear dependence with *T*_2_^C^ (Figure 2C).

**Figure 2.**
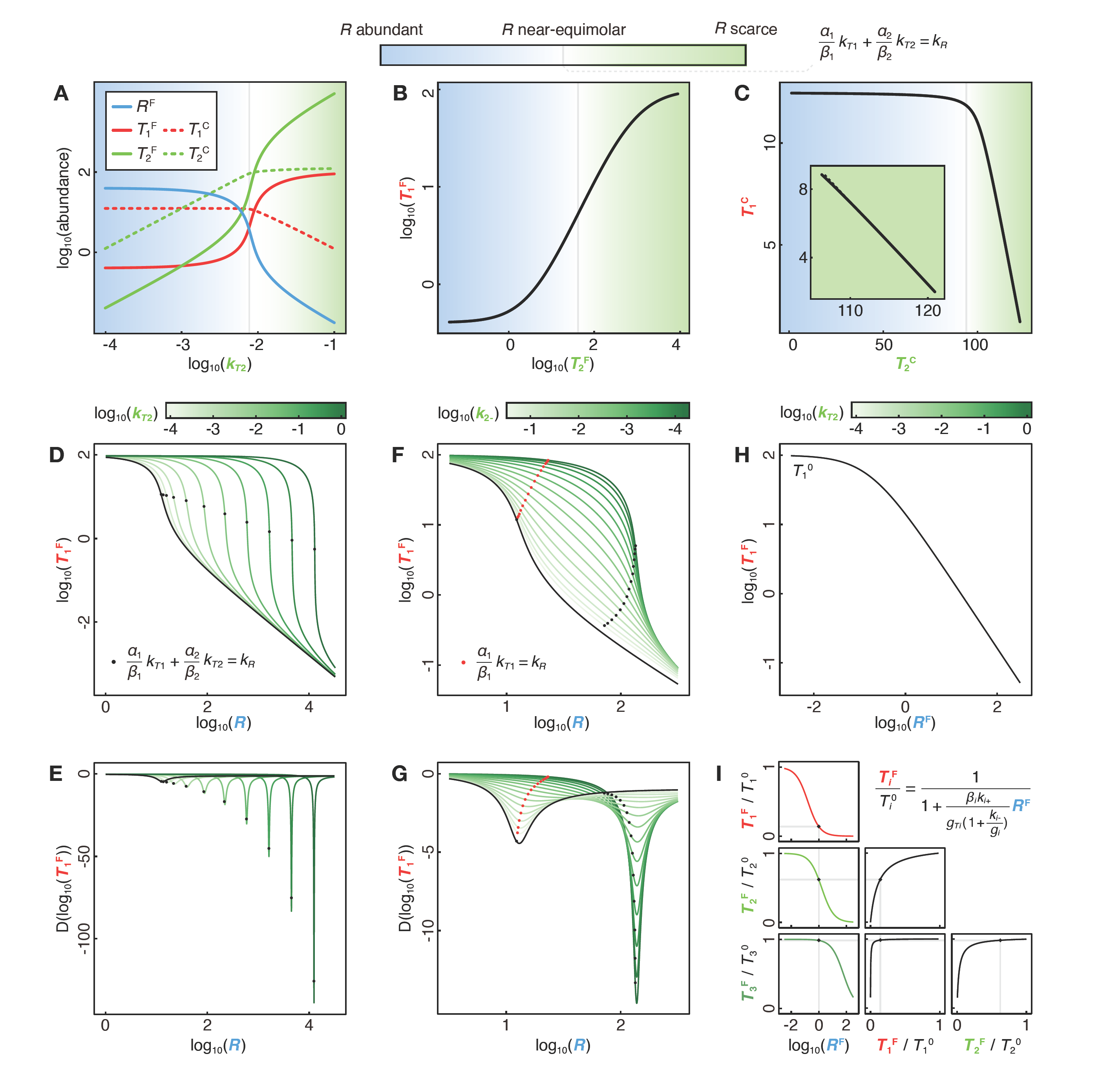
Steady state behaviors of competition systems. (**A-C**) Regimes of competition systems. (**A**) Abundances changes of each component with the increment of *T*_2_’s production rate (*k*_*T*2_). (**B**) Abundance of *T*_1_^F^ as a function of that of *T*_2_^F^ (**C**) Abundance of *T*_1_^C^ as a function of that of *T*_1_^C^. Blue, white and green areas represent *“R* abundant”, “R near-equimolar” and “R scarce” regime respectively. Grey lines represent the approximate threshold (SI Materials and Methods).(**D-G**) Dose-response curves modulated by competition. (**D-E**) *R*-*T*_1_^F^ dose-response curves (**D**) and their derivatives (**E**) with different *T*_2_’s production rate (*k*_*T*_). (**F-G**) R- *T*_1_^F^ dose-response curves (**F**) and their derivatives (**G**) with different *T*_2_^C^’s dissociation rate (*k*_2-_). *R* represents the total abundance of regulator (*R*^F^+*T*_1_^C^+*T*_2_^C^). Black lines represent the dose response curve without *T*_2_ (*k*_*T*2_=0).(**H**) R^F^-*T*_1_ dose-response curves with different *k*_*T*2_. *T*_1_^0^ represents the abundance of *T*_1_^F^ without *R*. Black line represents the dose response curve without *T*_2_ (*k*_*T*2_=0). All the curves with different *k*_*T*2_ are exactly overlapped.(**I**) Repression folds of all targets are determined by the same *R*^F^ abundance in a multitarget repression system.

In the case of ceRNA regulation, where miRNA is a repressor, target activity can be reflected by the free mRNA level. Increments of ceRNA_2_ (*T*_2_^F^) can raise free ceRNA1 (*T*1^F^) level indirectly by sequestering shared miRNAs. Such derepression caused by ceRNA effect is negligible when the level of ceRNA2 is far less than that of miRNA (in the *“R* abundant” regime), but becomes detectable when the level of ceRNA2 is comparable to that of miRNA (in the “R near-equimolar” regime) (Ala et al., 2013; Yuan et al., 2015). In contrast, when the regulator is an activator, target activity can be represented by the level of complexes. Recently a phenomenon called “isocost line” behavior, originally studied in economics, was also found in synthetic biological systems (Carbonell-Ballestero et al., 2016; Gyorgy et al., 2015) that the expressions of two fluorescent proteins in *E. coli* displayed negative linear dependence, which was caused by competition for the transcription and translation resources (acting as activator) by the two synthetic genes. Due to the high expression level of these genes, the system was always restricted to the “R scarce” regime, thus showed negative linear dependence.

In summary, threshold behavior and negative linear dependence are two aspects generated by the same competition motif. The threshold behavior is observed when the regulator is a repressor and the system transfers from the “R abundant” to the “R near-equimolar” regime; while the negative linear dependence occurs when the regulator is an activator and the system is restricted to the “R scarce” regime.

### Competition can shape dose-response curve

How does competition modulate the response of target to varying levels of a regulator? The dose-response curve, which quantitatively describes the magnitude of such responses, was systematically analyzed. Firstly, the dose-response curve of free *T*_1_ (*T*1^F^) level to the total regulator (**R**) level without competition effect (without *T*_2_) was calculated as the baseline. As expected (Buchler & Louis, 2008), *T*1^F^ was not sensitive to the regulator changes in the “R scarce” regime, but became sensitive in the “R near-equimolar” regime, thus forming some “threshold behavior” (black line in Figure 2D-E). Then we analyzed how the molecular levels and the kinetic parameters of the competitor *T*_2_ might influence the shape of the *R*-*T*1^F^ dose-response curve. We first considered the case that *T*_1_ and *T*_2_ have the same kinetic parameters to bind R. Increments of *T*2 production could elevate the maximum sensitivity to enhance the threshold behavior, and shift the position of the maximum sensitivity to a higher *R* level in the new “R near-equimolar” regime (Figure 2D-E). We next fixed *T*_2_’s production rate and analyzed the influence of other kinetic parameters. The relative binding affinity was found as the key parameters to modulate the R-*T*1^F^ dose-response curve. If *T*_2_^c^ was formed slowly (small *k_2+_*) or dissociated rapidly (large *k_2-_*), *T*_2_ could hardly alter the *R-T*1^F^ response. Along with the increment of *T*_2_ binding affinity (increasing *k_2+_*) or decreasing *k_2-_*)), *T*_2_’s competition blunted the sensitivity in the R~*T*_1_ near-equimolar regime considering only *R* and *T*_1_, meanwhile enhanced the sensitivity in the R~*T*_1_+*T*_2_ near-equimolar regime in the presence of *T*_2_ (Figure 2F-G and S2C-E).

The model analysis is consistent with the experimental observations in diverse molecular competition scenarios reported previously. In the case of ceRNA (Figure 1C), the RNA competitors with comparable binding affinities can enhance the maximum sensitivity and shift their positions in the miRNA-target dose-response curve, and a higher competing RNA level can cause a stronger enhancement and shift (Yuan et al., 2015). Similarly, in the studies on the TF titration effect (Figure 1B), introducing high affinity competitive binding sites can greatly shift and sharpen the response of primary target gene expression to the TF (Brewster et al., 2014; Lee & Maheshri, 2012). In contrast, in the case of buffer solutions in chemistry, for example the ammonium buffer, the weak base NH_4_^+^ compete with H^+^ for OH^-^, and NH_4_^+^ has a much lower binding affinity with OH^-^ than H+ (Figure S1E). When a mild change of OH^-^ (e.g. adding moderate amounts of NaOH or HCl) is introduced into the solution, NH_4_^+^ can buffer the response of free H^+^ to OH^-^, thus keeping pH (potential of hydrogen) almost constant in a certain range (SI Material and Methods). In summary, introducing the competitors can shape the *R*-*T*_1_^F^ dose-response curve. A high affinity competitor can enhance the maximum sensitivity and shift its position to a higher *R* level; while a low affinity competitor may buffer the response. The extents of such modulations are dictated by the abundance of competitors.

However, it should be noticed that when it comes to the response curve of a free primary target to the level of a free regulator (*R^F^*-*T*_1_^F^), the curve was not influenced by the existence of competitor at all (Figure 2H). This is because, rather than the total regulator abundance, the free regulator abundance is the one effectively determines the kinetic reaction rate with each single target (Jens & Rajewsky, 2015). Thus, responses of two or more targets to the shared regulator are mutually independent given the level of R^F^, which provides an efficient way, by using *R*^F^ level as the medium, to analyze the relative regulatory efficiency among multi-targets (Yuan et al., 2016). Once given the dose-response of each component (*R*^F^-*T_i_*^F^, which could be separately measured or calculated) and the expected regulatory efficiency of a specific target, the level of all other targets could be immediately predicted because they are all exposed to the same free regulator level (Figure 2I, SI Material and Methods). Such property is especially important for designing synthetic circuits, where we know the characteristics of each single part and would like to predict the whole system’s behavior when putting them together. This property has been applied to siRNA design principle: by both *in silico* simulation and experimental validation, we found that the influence of a high off-target gene expression level could be compensated by introducing a suitable number of siRNAs, whereas off-target genes with strong binding affinity should be avoided (Yuan et al., 2015; Yuan et al., 2016). In summary, the dose-response to the free regulator level is not influenced by any competitors, therefore providing an efficient way to extract the relative response relations in multi-target networks.

### Competition can delay or accelerate dynamic response

How does the existence of competitors influence the dynamic behavior of the system in response to a time-varying regulator? To answer this question, we simulated the response of a switching system with regulator level changing between “ON” and “OFF” states (Figure 3A). On the rising edge of R’s change, the existence of *T*_2_’s competition always delays the response of both *T*_1_^F^ and *T*_1_^C^, because it can sequester *R* from binding with *T*_1_ and may cause additional *R* loss via *T*_2_^C^ degradation, both of which resist the increment of available *R* to regulate *T*_1_. However, on the falling edge, competing can either accelerate or delay the response depending on the kinetic parameters (Figure 3B-C and S3A-F, SI Material and Methods). On the one hand, *T*_2_^C^ dissociation could compensate R’s decrease, but on the other hand, *T*_2_^c^ degradation may cause *R* loss, and these two opposing effects can dominate the final modulation of the dynamic response. *T*_2_ with a large complex degradation rate (*g*_2_) and a large loss rate (*α*_2_) could lead to a quick response by mediating more *R* loss (Figure 3B); while *T*_2_ with different binding affinities could either accelerate or delay the response under different parameter settings (Figure 3C and S3C-F), because *T*_2_ with a strong binding affinity can enhance both *R* compensation and *R* loss via *T*_2_^C^ degradation at the same time.

**Figure 3.**
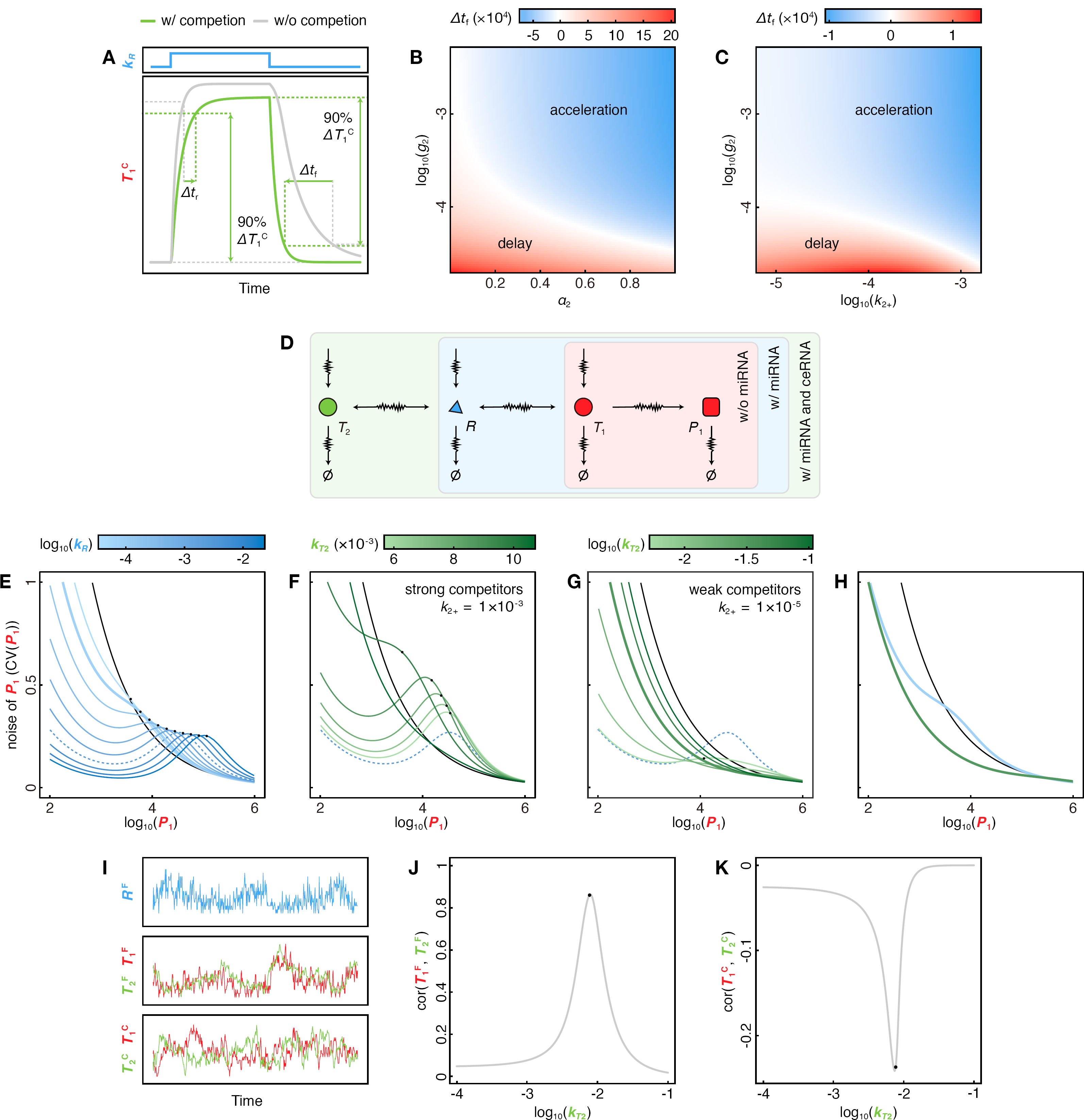
Dynamic properties of competition systems.(**A**) Quantitative measurements of response time. Δ*t_r_* and Δ*t_f_* represent the alteration of response time on the rising and falling edge of *R*’s change respectively. Here response time is defined as the time taken by *T*_1_^C^ level to change from 0% to 90% between its initial and final steady states.(**B-C**) Heatmaps of Δtf under different *α*_2_ and *g*_2_ (**B**), or *k*_2+_ and *g*_2_ (**C**).(**D**) Schematic diagram of the target expression noise in the miRNA-target competition scenario.(**E-H**) Modification of target expression noise by competition. (**E**) Product expression noise (CV(*P*_1_)) with different *R*’s production rates (*k_R_*). (**F**) CV(*P*_1_) with different T_2_’s production rates (*k*_*T*2_) where *T*_2_ acts as a strong competitor. (**G**) CV(*P*_1_) with different *kn* where *T*_2_ acts as a weak competitor. (**H**) Comparison of CV(*P*_1_) with or without competition. Here miRNA-RNA competing system is taken as an example. Black lines represent system without R. Dashed blue lines are highlighted as the basal lines in (**F**) and (**G**). The thick blue and green lines in (**H**) are taken from (**E**) and (**G**) respectively. Black dots represent the approximate threshold (there are no black dots on some curves because *kr2* is too large to form the threshold).(**I-K**) Correlated fluctuations introduced by competition. (**I**) Stochastic simulations of each component’s abundance in competition motif. (**J-K**) Correlations of *T*_1_^F^ and *T*_2_^F^ (**J**), or *T*_1_^C^ and *T*_2_^C^ (**K**) changing with *T*_2_’s production rate (*k*_*T*2_). Black dots represent the approximate threshold.

Recently, it has been experimentally observed that the competition for LacI binding in *E. coli* delayed the rising edge response, but accelerated the falling edge response because of the loss of the regulator binding with targets through degradation and dilution (large α_2_) (Jayanthi et al., 2013). On the contrary, the existence of competitive binding sites for transcription factor SKN7m in *S. cerevisiae* was found to delay the response of the primary target on both the rising and the falling edges (Mishra et al., 2014), which implied that the regulator might be protected from degradation when binding with targets (g_2_ is small) (Burger et al., 2010; Jayanthi et al., 2013). In summary, competition can modulate the dynamic response of some targets to their upstream regulators. This may implicate a general parameter tuning method to adjust the response dynamics in the presence of the competitors.

### Competition can modify target expression noise level

Competition can modulate the sensitivity and the speed of a target response to a changing regulator, both of which are highly relevant to target fluctuation (Blake et al., 2003; Chen et al., 2013). A natural question is how the existence of competitors may influence noise in the system? Here we took miRNA regulation as an example to analyze the noise level of protein products (Figure 3D, SI Material and Methods). In systems without *R* and *T*_2_, *T*_1_ expression noise is derived from fluctuations in transcription, translation and degradation, and the coefficient of variance (CV) of *T*_1_ gene expression approaches the “power law”, as expected by the “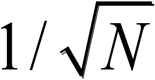 rule” proposed by Schrödinger (Schrödinger, 1944). The introduction of *R* (miRNA) as repressor can decrease the noise of lowly expressed genes, meanwhile generate a noise peak in the “R near-equimolar” regime for highly expressed genes (Figure 3E), consistent with previous studies (Bosia et al., 2017; Schmiedel et al., 2015).

Theoretical results indicated that the competition effect of *T*_2_ could modify *T*_1_ expression noise significantly. As expected, introducing *T*_2_ weakens *R*’s ability to suppress *T*_1_, thus may impair the noise reduction in the low expression zone. Interestingly, in the high expression zone of *T*_1_, *T*_2_ with strong binding affinity with *R* may elevate *T*_1_ noise level (Figure 3F); while *T*_2_ with weak binding affinity may substantially depress *T*_1_ noise level (Figure 3G). Therefore, comparing with the one-regulator-one-target scenario, introducing higher level of miRNAs and compensable weak competitors could reduce target expression noise at the low expression zone and suppress the noise peak introduced by miRNA at the high expression zone at the same time, thus could repress gene expression noise in a wide range (Figure 3H). In summary, competition effects may modulate gene expression noise level, and in particular, abundant weak competitors have the capability to buffer gene expression noise globally (Figure S3G-J).

### Competition can introduce correlated fluctuation between targets

Competition can not only modify the strength of target fluctuation, but also couple fluctuations between these targets (Figure 3I). Dynamic analysis of the model’s behavior around steady state with different molecular environments predicted that the free *T*_1_ (*T*_1_^F^) and *T*_2_ (*T*_2_^F^) are positively correlated (Figure 3J), while the competitor complexes (*T*_1_^C^ and *T*_2_^C^) are negatively correlated (Figure 3K). The correlation strengths in both cases are maximized in the “R near-equimolar” regime, and gradually decrease with the system away from the regime.

This phenomenon has been predicted as the “correlation resonance” by some previous theoretical analysis on gene translation (Mather et al., 2013) and protein degradation (Cookson et al., 2011; Mather et al., 2010). Two kinds of proteins (*T*_1_^F^ and *T*_2_^F^) competing for degradation enzyme ClpXP (**R**) showed positive correlated fluctuation, which reached the maximum when the sum of two protein production rates approached to the ClpXP’s processing capacity (Cookson et al., 2011; Mather et al., 2010). Another theoretical analysis showed that in translation process, fluctuations of mRNA-ribosome complexes (*T*_1_^C^ and *T*_2_^C^) were negatively correlated (Mather et al., 2013). In summary, competition can introduce negatively correlated fluctuation between free targets and positively correlated fluctuation between complexes, and both of their strength reach the maximum in the “R near-equimolar” regime.

### Regulator allocation to multiple targets

Regulators often bind more than two target species simultaneously. How will regulator be allocated to multiple target species? A system with multiple targets competing for the same regulator can be described by the set of allocation equations (Figure 4A), where the proportion of the regulator occupied by a certain target in steady state is mainly determined by this target’s abundance and its capabilities to bind to (and hence to consume) the regulator (SI Material and Methods). It was noticed that, the form of the regulator allocation equation is analogous to Kirchhoff’s laws in current divider circuits, where *R*’s production rate is analogous to the total current, the capability of *T*_i_^C^ to consume *R* is analogous to the *i*th branch current, and the capability of *T*_i_^F^ to occupy *R* is analogous to the ith branch conductance (the reciprocal of resistance) (Figure 4B). Therefore, electronic circuits and biological systems with competition may exhibit similar properties, such as the “negative linear dependence” behavior when resources are insufficient (in the “R scarce” regime) (Carbonell-Ballestero et al., 2016).

**Figure 4.**
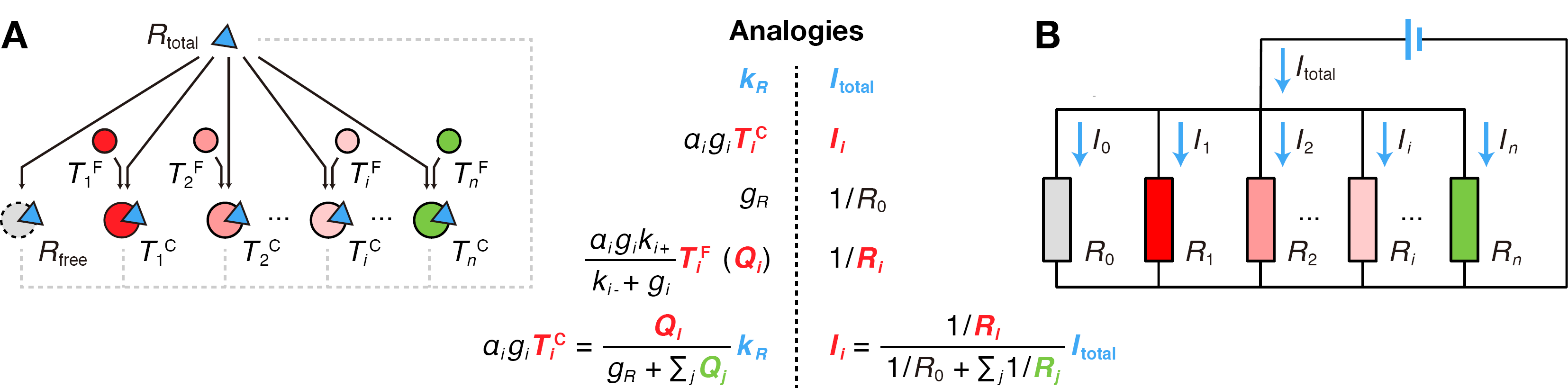
Regulator allocation for multi-target competition.(**A**) Regulator allocation equations and schematic graph representation. *R*_total_ represent the total abundance of regulator, including free regulator and regulator in complexes.(**B**) Kirchhoff’s laws in current divider circuits.

Such allocation equations have displayed in diverse mathematical models, such as the reaction rates of product formation in enzymatic reactions when multiple substrates competing for the same catalytic enzyme under the Michaelis-Menten kinetics (Chou & Talaly, 1977), and the probabilities of promoter-TF binding when multiple promoters competing for the same TF under the thermodynamic model (Bintu et al., 2005). Meanwhile, this property has helped quantify the allocations of the transcription or the translation resources for synthetic gene circuits (Carbonell-Ballestero et al., 2016; Qian et al., 2017). We also applied such property to predict the miRNA occupancy on each target site in a specific cell type with the miRNA and the target RNA expression levels, and significantly improved the accuracy of the miRNA target prediction (Xie et al., 2014). Those miRNAs with significant occupancy changes during tumorigenesis could serve as potent biomarkers in addition to differentially expressed miRNAs.

## Discussion

Competition for limited resources is ubiquitous throughout diverse molecular reactions in both natural and synthetic biological systems. Using a coarse-gained mathematical model, we systematically analyzed the steady-state behavior and the dynamic properties of various competition network motifs, from the view of indirect regulations among the competitors as well as the effects of the competitors on the regulator-target response (Table 1). It should be noticed that, most of the mentioned properties are connected with the concept of the *regimes* determined by the regulator-target relative abundance (Figure 2A-C): threshold behavior occurs when system transfers from the *“R* abundant” to the *“R* near-equimolar” regime, and linear negative dependence happens when system is in the “R scarce” regime; while the sensitivity of the dose-response curve, the correlated fluctuation, and the noise of the target level are all maximized in the “R near-equimolar” regime.

**Table 1.** Properties of regulation by competition

|  | **Regulation between targets** | **Influences on**<br>**regulator-target response** |
| --- | --- | --- |
| **Steady-state**<br>**behavior** | Threshold behavior<br>Negative linear dependence<br>Regulator allocation | Shaping dose-response curves |
| **Dynamic**<br>**responses** | Correlated fluctuation | Response time modulation<br>Noise modification |

Competition motif is a common network component. It seldom functions as an isolated module in real-world biological systems, but often interacts with other components to form complex networks. For example, simulation analysis on ceRNA regulation suggested that additional targets and regulators connected with different topology could enhance or weaken the ceRNA effect (Ala et al., 2013). Theoretical analysis predicted that competition for degradation enzyme could either promote or suppress the robustness of biological oscillating circuit with different topological structures (Rondelez, 2012). In addition, competition motif could perform a variety of functions by combining with other network motifs. For example, cooperating with the positive feedback motif, competition can generate the winner-take-all (WTA) behavior (Kim et al., 2004), which have been applied to design *in vitro* molecular circuits for supervised learning and pattern classification using DNA strand displacement (Genot et al., 2013; Lakin & Stefanovic, 2016).

The unified competition model gives inspirations for transferring knowledge among different molecular scenarios, since similar molecular network topology may perform similar functions. For example, the case that ceRNA competition can sharpen the dose-response curve of miRNA regulation (Yuan et al., 2015) is quite similar to that observed for TF titration effect (Brewster et al., 2014). Such generality and feasibility give us confidence to make new predictions based on the competition model. For instance, the properties of pH buffer solutions demonstrated that some weak competitors could desensitize the response of the primary target to the regulator, which implies the potential role of many competitors as noise buffer. Functions of numerous miRNA target sites have long been a mystery that each miRNA species in mammalian cell could bind to hundreds target RNA species, but only a small portion of the targets with multiple high affinity binding sites could be moderately repressed (rarely exceeds 2-folds). That is to say, in most cases, miRNA binding are not functioned as intensive repression (Seitz, 2009). Why are there so many evolutionary conserved miRNAs and potential targets if this is an inefficient regulatory mechanism? The competition model provides a possible explanation that such widespread miRNA competitors with low binding affinity could buffer noise and stabilize gene expression.

Competition effect is one of the major challenges for circuits design in synthetic biology. Synthetic gene expression can lead to intracellular resource reallocation, which may affect the performance of both exogenous gene circuits and host gene networks simultaneously. It may change the network structure of the original designed circuits by introducing a hidden layer of regulation, making it difficult to predict the whole circuit’s behavior based on the characteristic of each individual component. For example, competition for cellular resources may reshape the response of genetic activation cascades in *E. coli* (Qian et al., 2017), and multiple downstream genes competing for upstream signal molecules may accentuate the “retroactivity” (Brophy & Voigt, 2014). It has been found that the induction strength of the synthetic gene oscillator could influence the growth rate of host cell, the expression of endogenous genes, and the performance of the oscillator, such as amplification and period (Weisse et al., 2015). On the other hand, interestingly, using competition effect properly to rebalance synthetic circuits’ relation to the host cell is emerging as an effective way to refine circuits performance. For example, the robustness of the synthetic oscillator can be greatly improved by introducing competing binding sites for TF LacI to sharpen target gene dose response curves and suppress gene expression noise (Potvin-Trottier et al., 2016). Models incorporating circuit-host competition effects can predict synthetic gene behaviors better (Liao et al., 2017). Reallocating the cellular translational resources by introducing the endoribonuclease MazF circuit can significantly enhance exogenous enzyme expression to promote metabolite production (Venturelli et al., 2017). Utilizing synthetic miRNA and its competitive binding RNA sponges, a RNA-based AND gate circuit was designed for selectively triggering T cell-mediated killing of cancer cells (Nissim et al., 2017).

As discussed in this paper, competition of molecules matters in diverse biological processes, not only convoluting regulations in cell, but also introducing plentiful functions. The concept of competition motifs and its coarse-gained model may provide a unified insight to understand diverse molecular competition phenomena, and modulate biological networks by coupling or decoupling components on the hidden layer.

## Materials and Methods

Detailed information about mathematical derivations and simulations is available in SI Materials and Methods. Parameters for simulations are shown in Table S1.

## Acknowledgements

This work has been supported by the National Science Foundation of China Grants (No. 61773230, 31371341, 61721003, 91730301, 31671384, 91729301), National Basic Research Program of China (2017YFA0505503), Initiative Scientific Research Program (No. 20141081175) and Cross-discipline Foundation of Tsinghua University, and the Open Research Fund of State Key Laboratory of Bioelectronics Southeast University.

## SI Materials and Methods

### 1. A unified coarse-gained competition motif model

Parameters involved in the competition motif model (Figure 1F) where two target molecule species (target#1 and #2, *T*_1_ and *T*_2_) competitively bind with a shared regulatory molecule species (regulator, *R*) are described as follows. In general, *T*_1_, *T*_2_ or *R* is produced with a rate of *k_T1_*, *k*_*T*2_ or *k_R_*, respectively. Free *T*_1_ 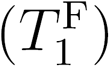, *T*_2_ 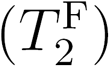 or *R* (*R*^F^) degrades at a rate of *g*_*T*1_, *g*_*T*2_ or *g*_*R*_. 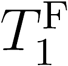 or 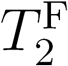 binds to *R*^F^ to form target-regulator complex 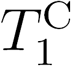 or 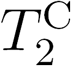 at a rate of *k*_1+_ or *k*_2+_, and 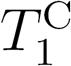 or 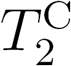 dissociates into *R*^F^ and 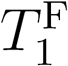 or 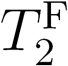 at a rate of *k*_1-_ or *k*_2-_. 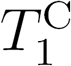 or 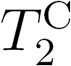 degrades at a rate of *g*_1_ or *g*_2_. Regulators on the complex degrade with the possibility of *α*_1_ or *α*_2_, and targets on the complex degrade with the possibility of *β*_1_ or *β*_2_, thus regulator would recycle from 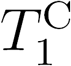 or 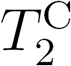 with the possibility of 1 − *α*_1_ or 1 − *α*_2_, target would recycle from 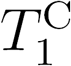 or 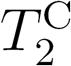 with the possibility of 1 − *β*_1_ or 1 − *β*_2_, and regulator and target would degrade together with the possibility of *α*_1_ + *β*_1_ − 1 or *α*_2_ β *β*_2_ − 1. When *R* is a repressor, 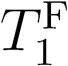 or 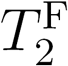 may generate production *P*_1_ or *P*_2_ at a rate of *k*_*P*1_ or *k*_*P*2_. In contrast, when *R* is an activator, 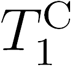 or 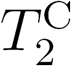 may generate production *P*_1_ or *P*_2_ at a rate of *k*_*P*1_ or *k*_*P*2_. *P*_1_ or *P*_2_ degrades at a rate of *g*_*P*1_ or *g*_*P*2_.

The competing model is described in the following differential equations:

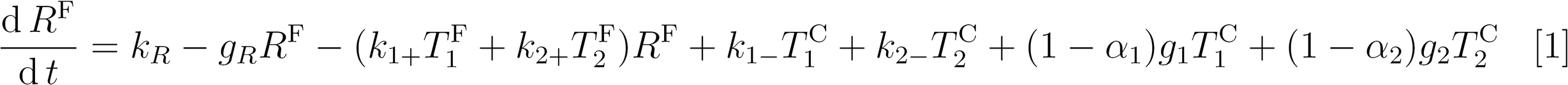

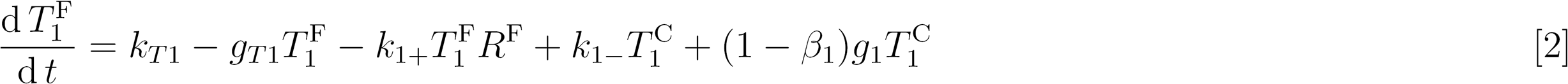

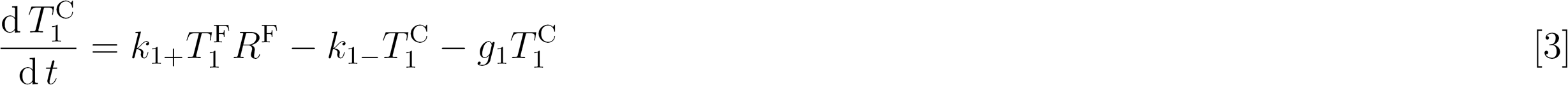

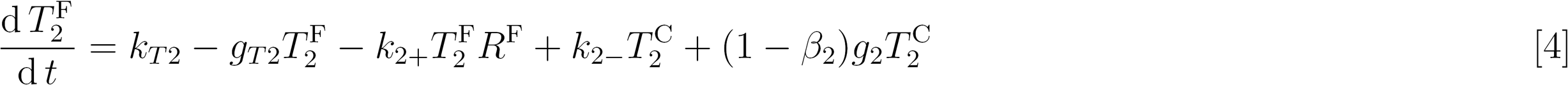

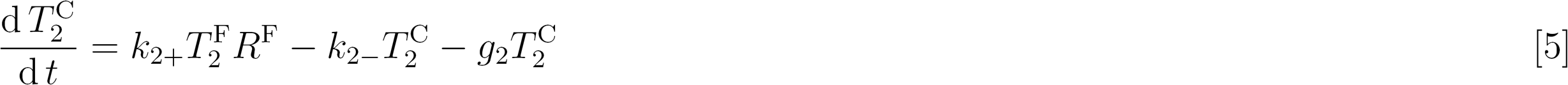

We used this model to describe competitions in various biological processes. In the competition for TF by DNA binding sites (Figure 1B), *T*_1_ and *T*_2_ represent TF binding sites on DNA and *R* represents TF. The production and degradation rates of DNA binding sites are set to zero because they are negligible. Complexes degrade with only TF loss (*α* ~ 1, *β* ~ 0). When *g*_1_ or *g*_2_ are set to zero, there is no TF loss. For TF as activator, DNA-TF complexes (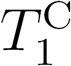 and 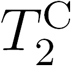) can be transcribed into RNA, while for TF as repressor, free DNAs (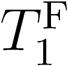 and 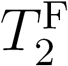) can be transcribed (Figure S1A). In the competition for miRNA by RNA molecules (Figure 1C), *T*_1_ and *T*_2_ represent two RNA molecule species and *R* represents miRNA. The loss of miRNA is relatively small so *β* is set to zero (Figure S1B) and as miRNA acts as a repressor, only free RNAs (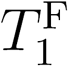 and 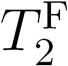) translate into proteins. In the case of ribosome allocation (Figure 1D), where *T*_1_ and *T*_2_ represent two RNA molecule species and *R* represents ribosome, *β* is also set to zero (Figure S1C). In protein degradation competition (Figure 1E), where *T*_1_ and *T*_2_ represent two protein molecule species and *R* represents the protein degradation machine, *β* is set to zero too (Figure S1D). The topology of miRNA-target competition, ribosome-mRNA competition and protein degradation competition are identical except that components generating further production are different.

### 2. Theoretically analysis for molecular environment determining shapes of the regulation between competitors

#### 2.1. Solving steady states

Eqs. 1-5 can be solved for steady state when giving all differentials as zero. By adding Eqs. 2 and 3, we get

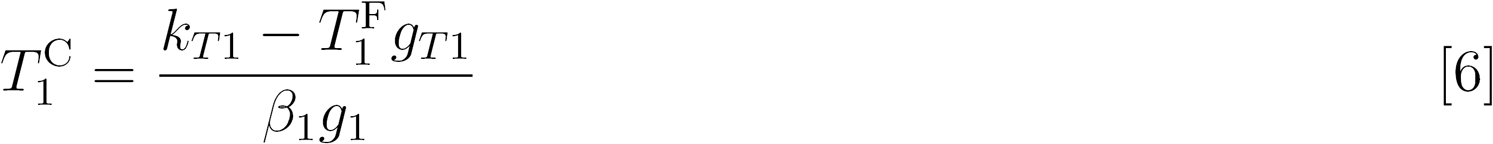

By adding Eqs. 1, 3 and 5, we get

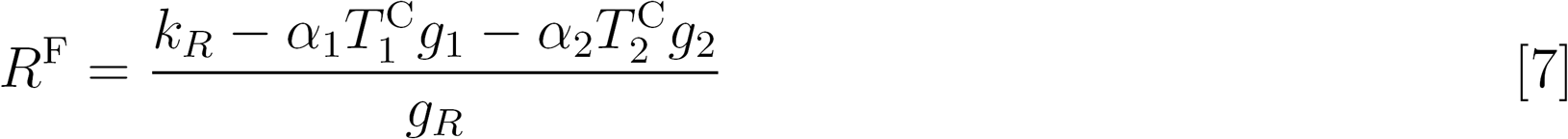

Combining Eqs. 6 and 7, we get

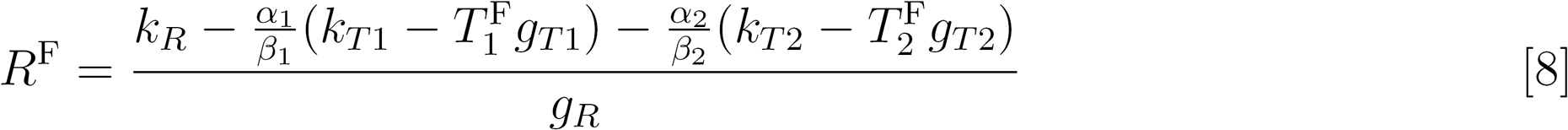

Substituting Eqs. 6 and 8 into Eq. 3, we get

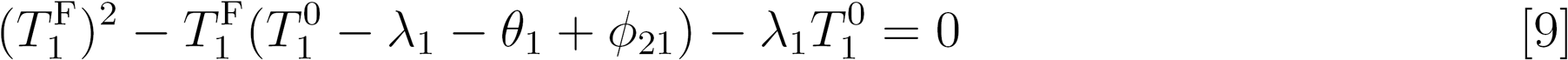

Where

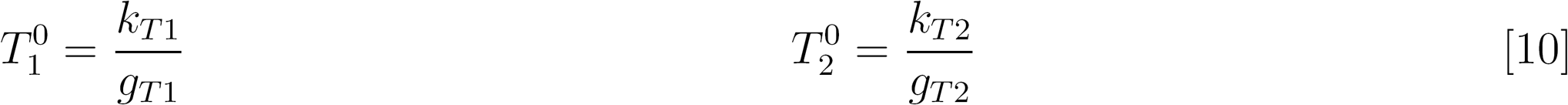

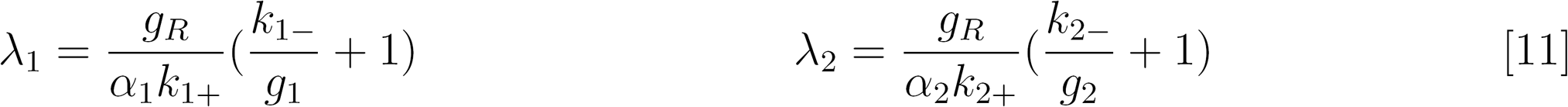

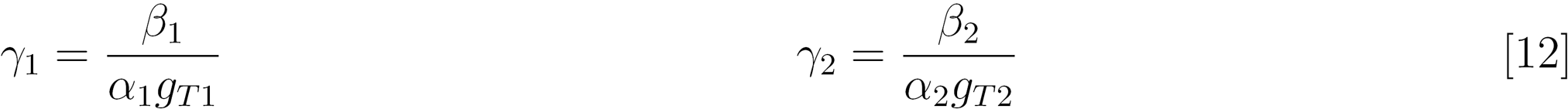

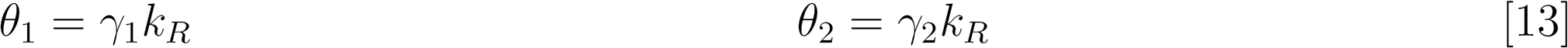

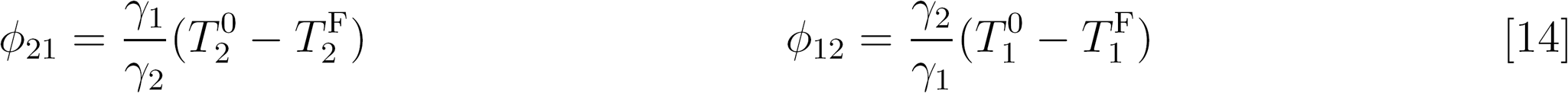

Parameters were lumped to represent certain physical meanings to simplify the result. 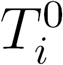 represents the free level of target *#i* (*T_i_*) without regulators. 1/*λ_i_* is proportional to *k*_*i*+_, and negatively correlated with *k*_*i*-_, thus could reflect the strength of binding affinity between *T_i_* and regulator. *θ* is proportional to *k_R_*, thus could reflect the level of regulator. *ϕ_ji_* exhibits the competing regulation effects by target *#j* upon to target *#i*.

Eq. 9 is a quadratic equation of 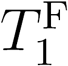. Thus, the steady state abundance of free targets can be expressed as

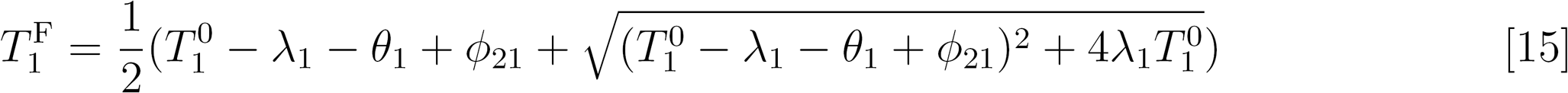

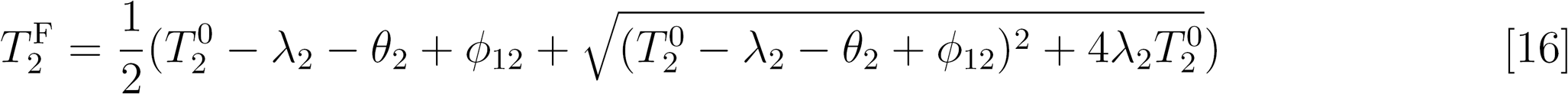

##### 2.2. Explanations on regimes and related phenomena

Assuming that the binding between targets and regulator is very strong, *λi* becomes negligible, thus Eq. 15 can be simplified as follows:

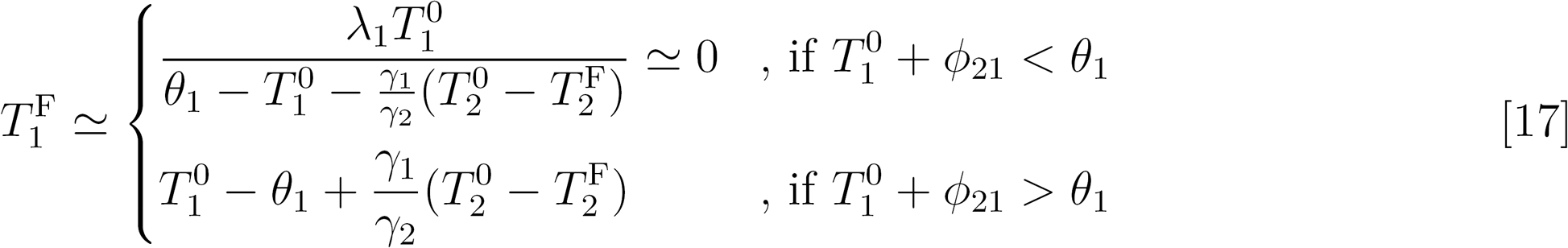

Meanwhile, the steady-state abundance of 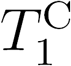 and 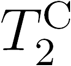 can be calculated from Eq. 6:

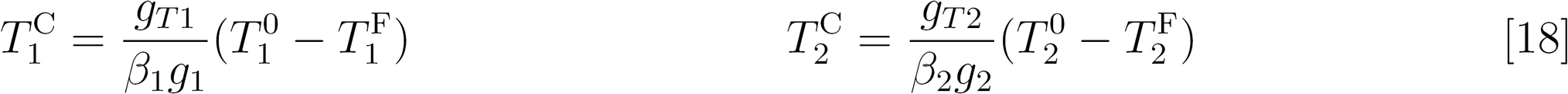

and can be simplified using Eq. 17:

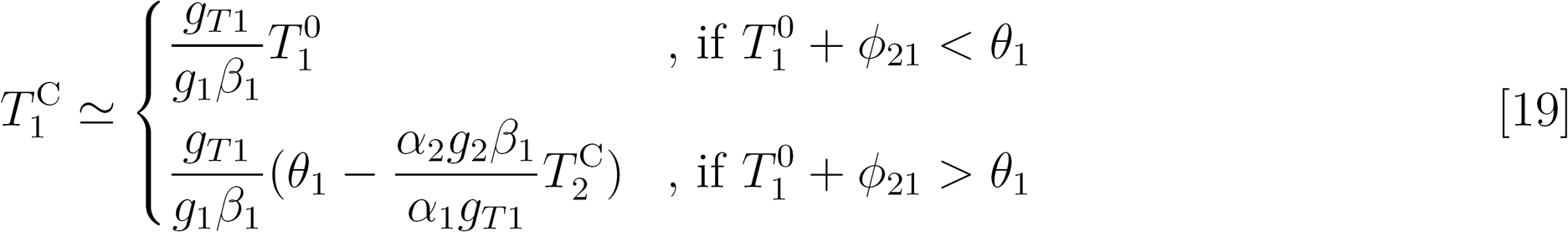

The turning point in Eqs. 17 and 19:

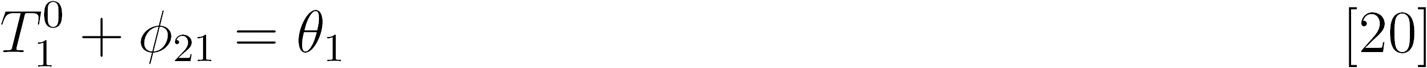

can be regarded as a threshold to distinguish regimes of the system: “*R* abundant” (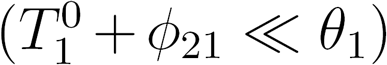, “*R* equimolar” 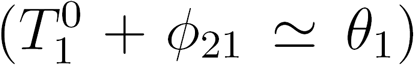 and “*R* scarce” 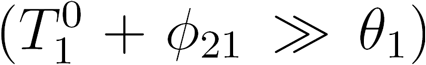. Eqs. 17 and 19 explain why the relationships between competitors are piecewise (Figure 2A-C). For 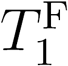 and 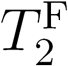, according to Eq. 17, in “*R* abundant” regime 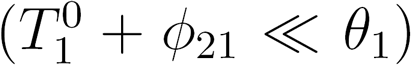, almost all targets bind with *R*, so the level of 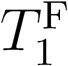 approaches to zero. In the contrary, in “*R* scarce” regime 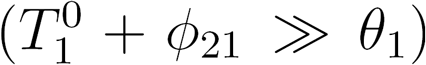, 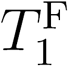 increases with the increment of 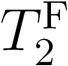, which is because when the production rate of *T*_2_ raises to sequester *R*, 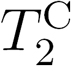 increases thus 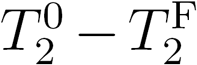 increases according to Eq. 18. Given the above, when the production rate of *T*_2_ increases to switch the system from *R* “abundant” regime to “*R* scarce” regime, the abundance of 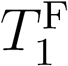 will exhibit a “threshold behavior” (Figure 2B).

Similarly, Eq. 19 suggested that the relationship between 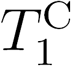 and 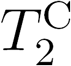 is piecewise linear. If *R* is abundant 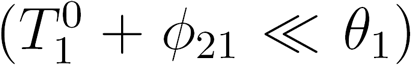, 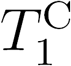 would keep substantially unchanged, while when *R* is scarce 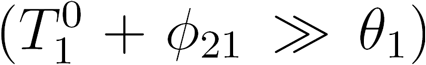, 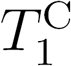 would decrease linearly with the increment of 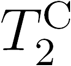, thus shows “negative linear dependence” (Figure 2C).

#### 2.3. Approximation of the regime threshold

The threshold (Eq. 20) can be approximated based on the strong binding assumption. It is equivalent to:

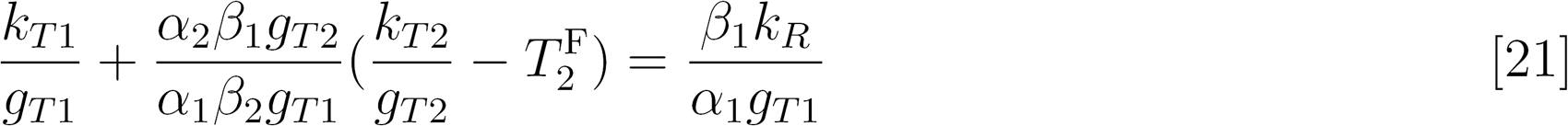

From Eq. 3, we get

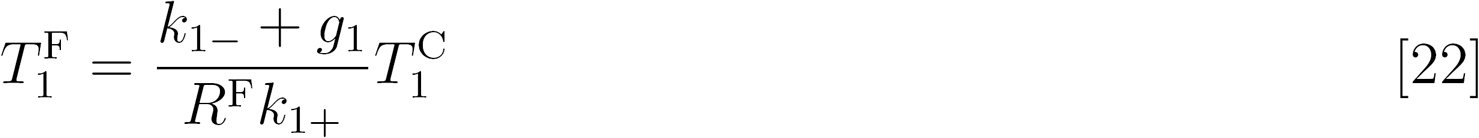

According to Eq. 17, before the system reaches the threshold 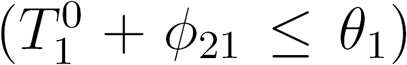 in the process of increment of production of *T*_2_, 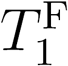 is much smaller than 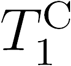 and approaches to zero, and so does 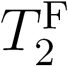. Thus, the threshold point 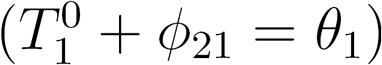 could be approximated from Eq. 21 as:

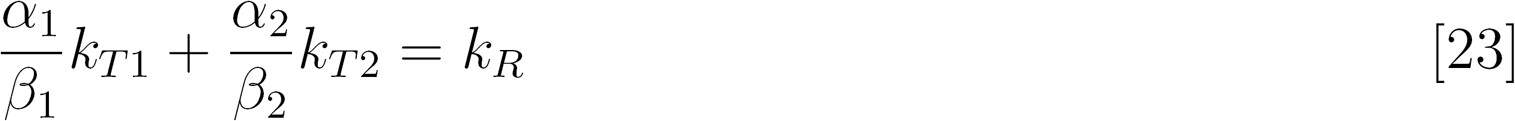

Eq. 23 gives an approximation of the threshold position to estimate the regime of a competing system roughly.

### 3. Competition can shape the regulator-target response curve

#### 3.1. How competition shapes regulator-target response curve

According to Eq. 9, there are

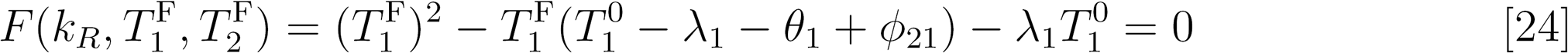

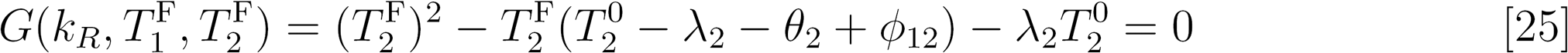

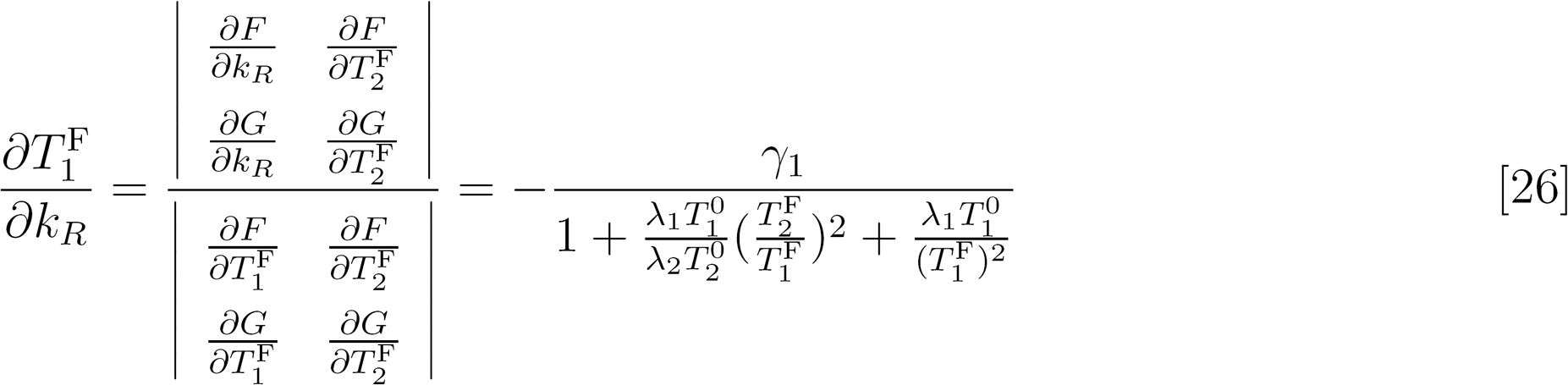

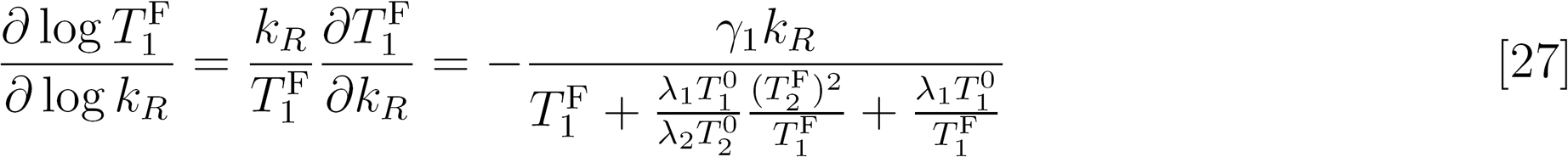

Eq. 27 describes the derivative of regulator-target response curve (Figure 2E and 2G).

Similarly, the buffer capacity, which quantifies the ability to resist pH changes in buffer solution, can be calculated as

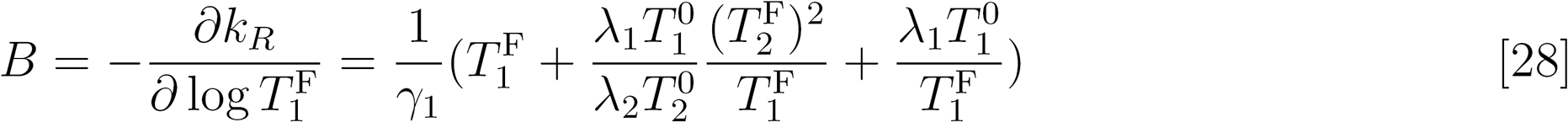

#### 3.2. Competition in buffer solution

For any buffer solution with a weak acid (HA) and its conjugate base (A^-^) or a weak base (BOH) and its conjugate acid (B^+^), there are

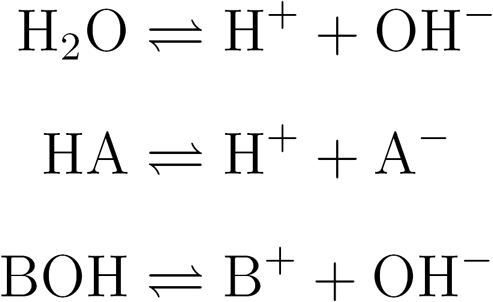

Here we take ammonium buffer solution (NH_3_. H_2_O and NH_4_Cl) as an example. In a aqueous solution with *a* mol/L NH_3_.H_2_O and *b* mol/L NH_4_Cl, there are

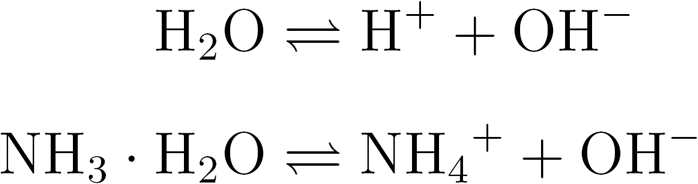

The equilibrium constants of these two reactions are

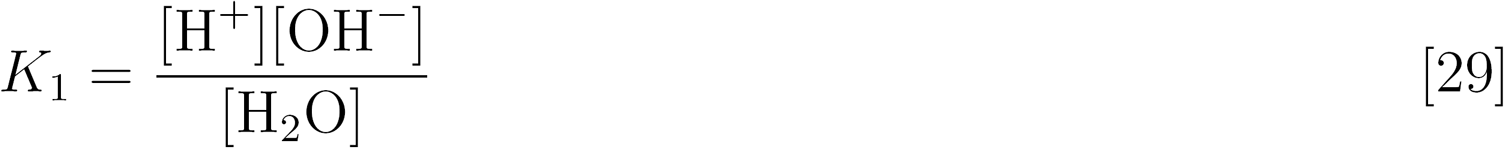

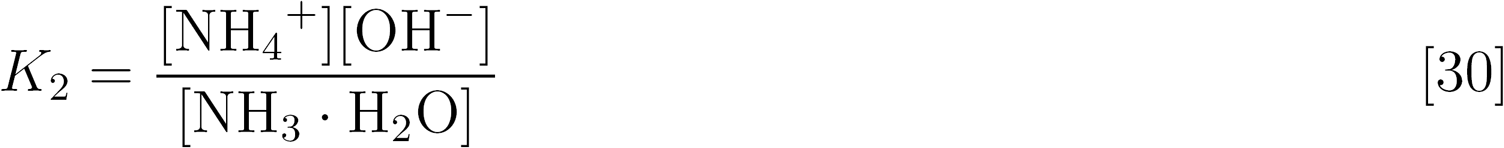

Because the concertation water of in an aqueous solution is almost invariant, the equilibrium constant of water (ion-product constant) is defined as

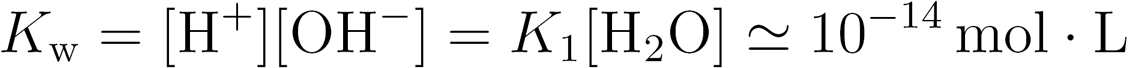

Here, we consider H^+^ (*T*_1_) and NH_4_+ (*T*_2_) competing for OH (*R*). Thus, Eqs. 29 and 30 is equivalent to

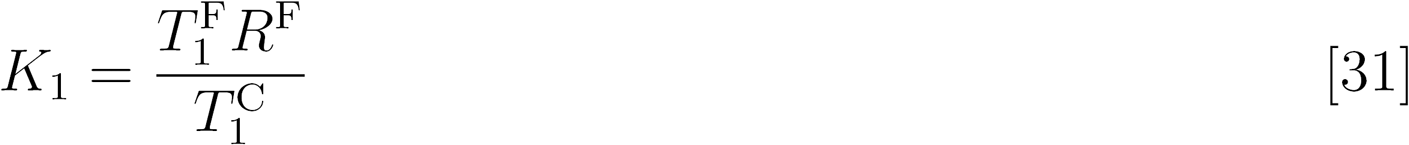

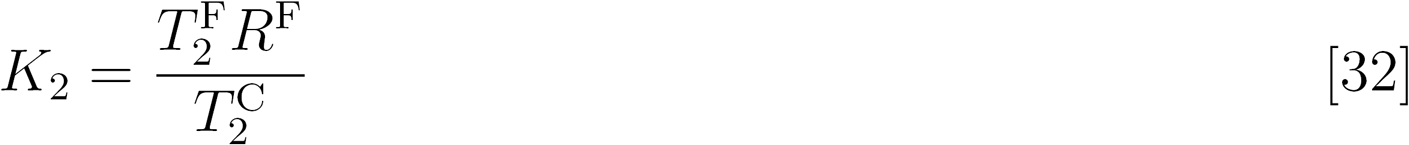

Meanwhile, because there are no production and degradation of any component, every substance is conserved as

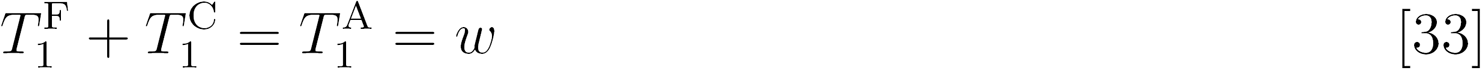

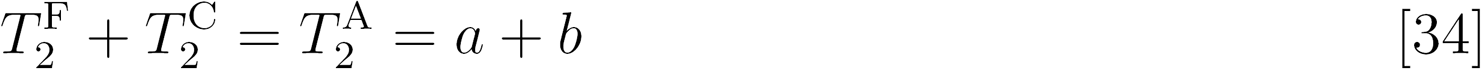

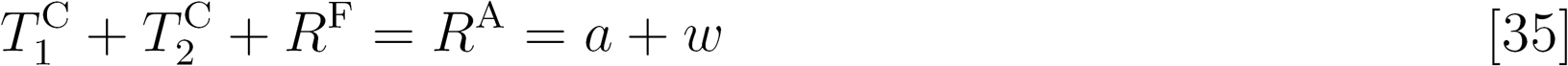

Combining Eqs. 33-35, we get

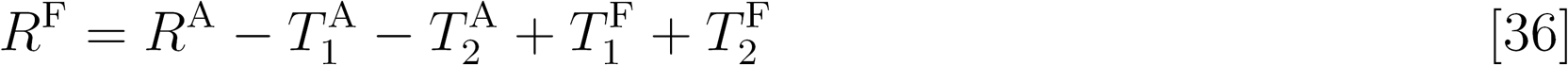

Combining Eqs. 31, 32 and 36, we get

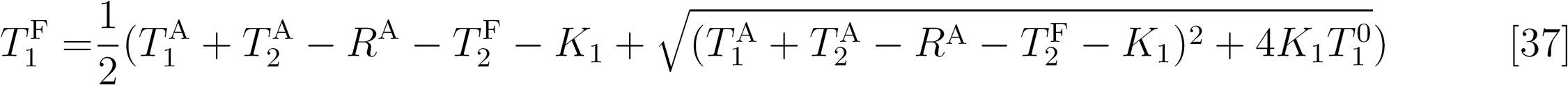

which is a degenerate form of Eq. 15, where

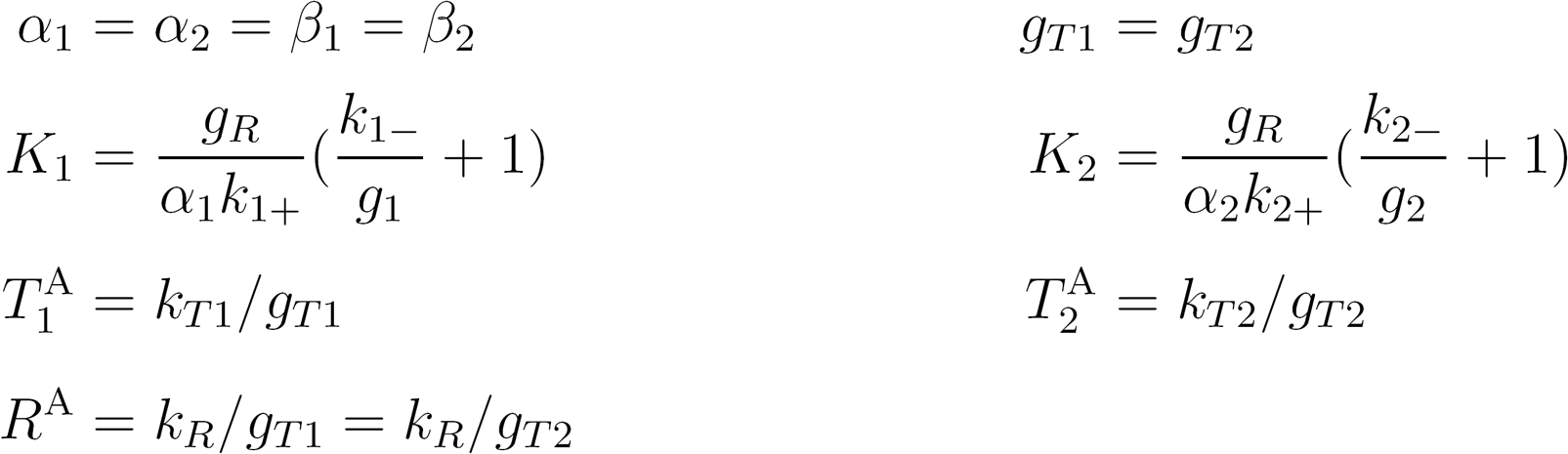

According to Eq. 28, the buffer capacity of this solution is

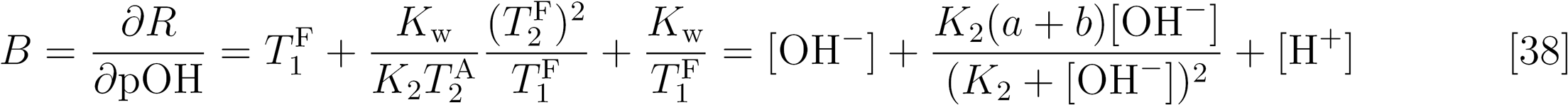

Eq. 38 indicates that when a mild change of OH^-^ is introduced to the solution, the buffer capacity guarantees the stable of pOH (and pH). More buffer substance (NH_4_+ and NH_3_.H_2_O, *a + b*) can lead to a larger buffer capacity, and the buffer capacity may maximize when pH = p*K*_2_.

#### 3.3. Dose-response curve of free target to free regulator

Substituting Eq. 22 into Eq. 2, we get

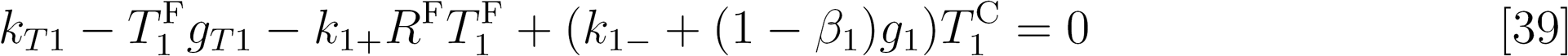

Thus,

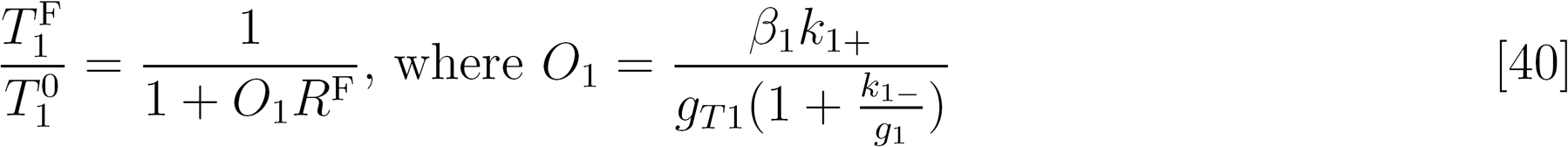

In system with *n* targets competing for same regulator, for the ith target (*i* =1, 2, …, *n*), this result can be extended as:

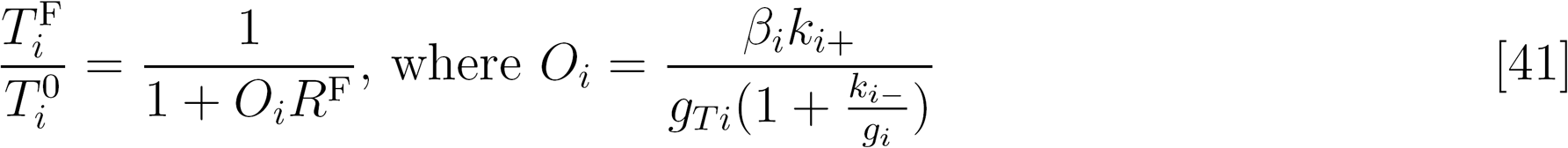

Similarly,

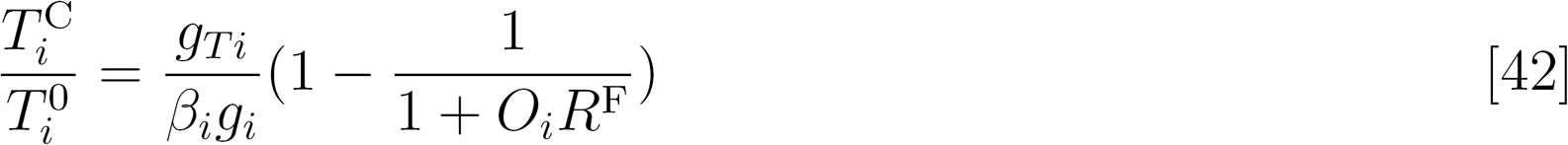

Eqs. 41 and 42 indicate that the level of 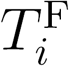 and 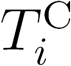 are determined only by the free level of R, and some chemical kinetic parameters of *T_i_* and *R*. In another wordsif two or more tagets compete for shared *R*, the relative abundances of each free target or complex are independent of other targets when giving the free level of *R*. In the siRNA design strategy (Yuan et al., 2015; Yuan et al., 2016),this property guarantees that no matter what expression of the off-target gene is (unless the expression is zero), the amount of free siRNA required to repress the target gene to a certain extent would always repress the off-target gene to a certain extent, which is determined by *O*_on_ and *O*_off_, as described in Eq. 41. When giving the expression of any target gene, siRNA could act as a medium to predict the expression of other target genes. This property also guides how to select suitable chemical reaction parameters: a good siRNA should have large *O*_on_ and small *O*_off_.

### 4. Competition can delay or accelerate dynamic response

When *R* level changes, comparisons of 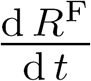 tells how competition affects the dynamic response speed of *T*_1_ with respect to *R*. According to Eq. 1 and 5,

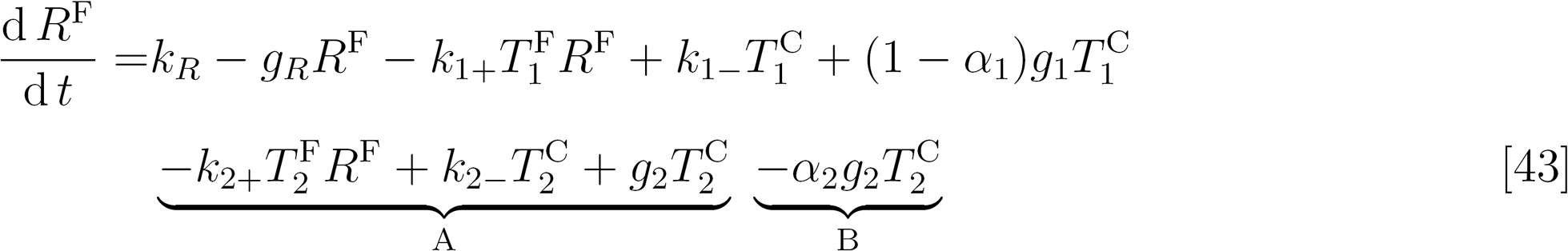

Item A equals to 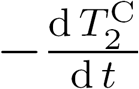, and indicates the ability of *T_2_* to sequester *R* when 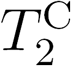 forms 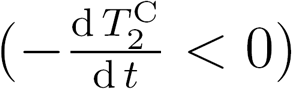, or release *R* when 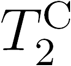 dissociates 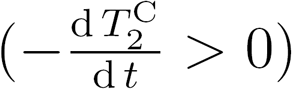. Item B indicates the level of *R* loss mediated by 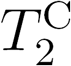 degradation.

On the rising edge of *R*, 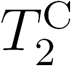 forms so item A < 0, meanwhile item B < 0 all the time, thus 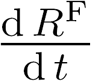 is smaller than non-competing system, leading to a slower response. On the falling edge of R, item A > 0 while item B < 0, thus the response speed depends on the relative magnitude of item A and B. As *g*_2_ or *α*_2_ increases, the absolute value of item B increases to alter the response from delay to acceleration. As 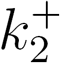 increases or 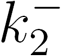 decreases, item A increases while the absolute value of item B also increases because 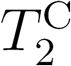 becomes larger, thus delay the response when there is no *R* loss mediated by 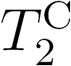 (*g*_2_ = 0, Figure S3D), alter the response from delay to acceleration when *g*_2_ is moderate (Figure S3E), or accelerate the response when *g*_2_ is large enough (Figure S3F).

### 5. Noise and correlated flucuation evaluation

The variances and co-variances of the molecular species in the system can be estimated with linear noise approximation. Fluctuation-dissipation theorem provides a general way to quantifies the fluctuations. The fluctuation-dissipation equation was solved numerically to calculate the covariance matrix *C*, the diagonal elements of which are the variance of corresponding entities, while the off-diagonal elements of which describe the co-variances between molecular species. The noise of a molecular species *i* is defined as coefficient of variation 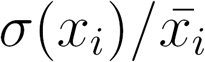 and the correlation between two molecular species *i* and *j* is defined as cov(*x*,*y*)/(*σ*(*x_i_*)*σ*(*x_j_*)), where *x_i_* and *x_j_* are random variables representing the abundance of molecular species *i* and *j*.

Taking miRNA competing system as an example (where miRNA acts as repressor), the vector of molecular number

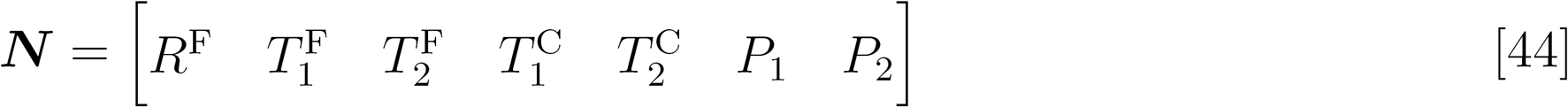

Transition rates vector is

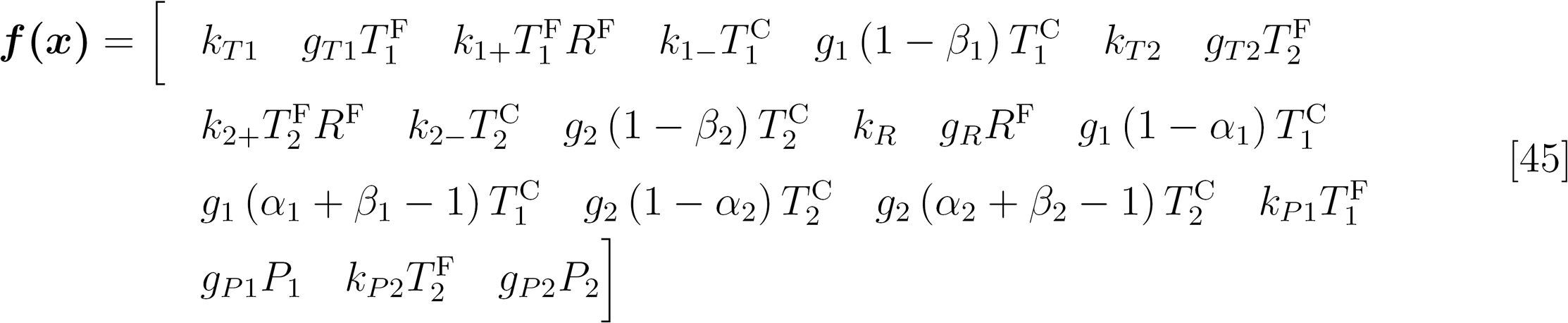

Stoichiometric matrix is

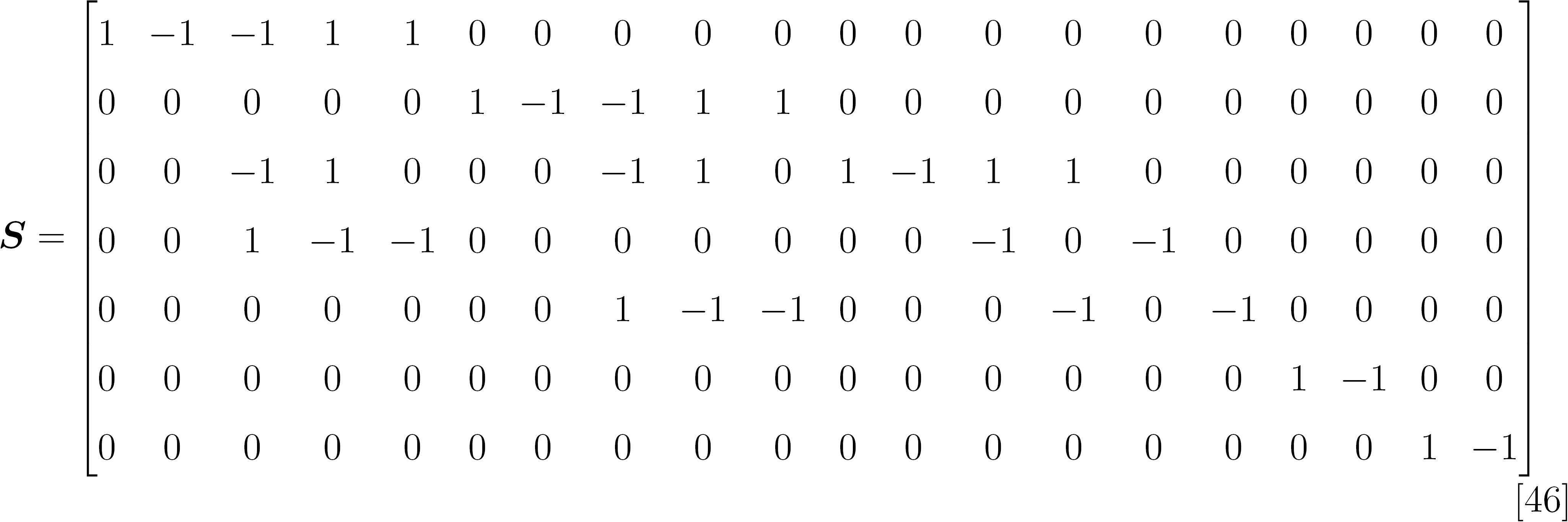

In the steady state, the rate equations can be linearized by the Jacobian matrix:

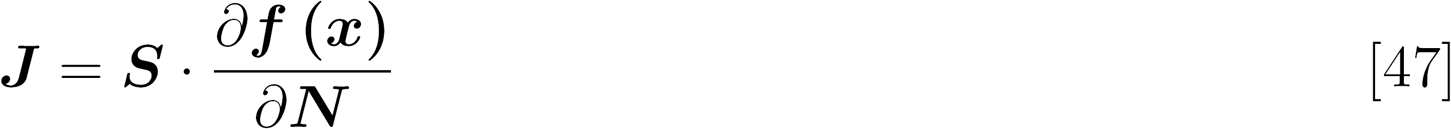

The diffusion matrix ***D*** is

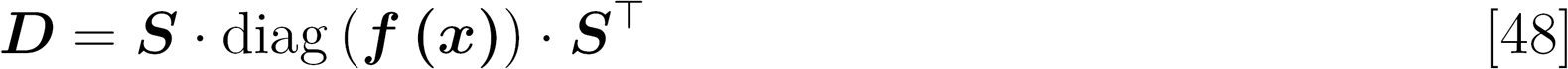

Therefore, the covariance matrix ***C*** can be calculate numerically by solving the fluctuation dissipation equation:

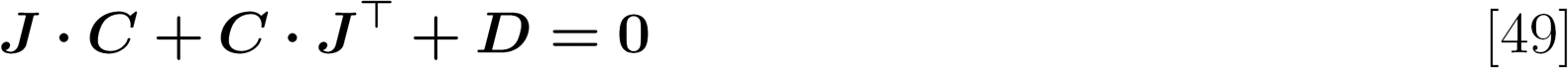

### 6. Regulator allocation to multiple targets

When there are *n* targets, similarly to Eqs. 1-5, there are

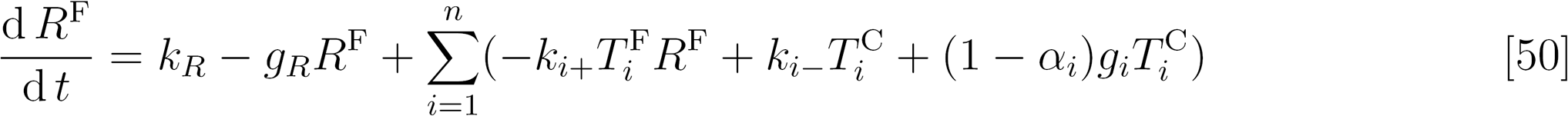

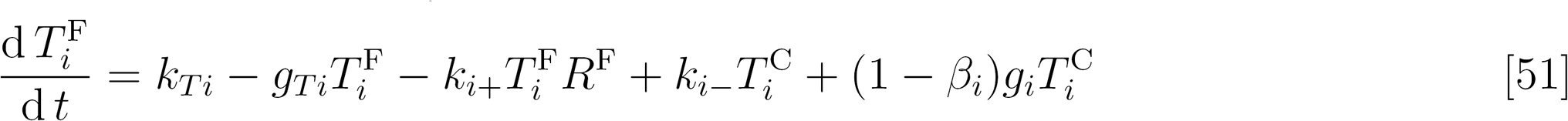

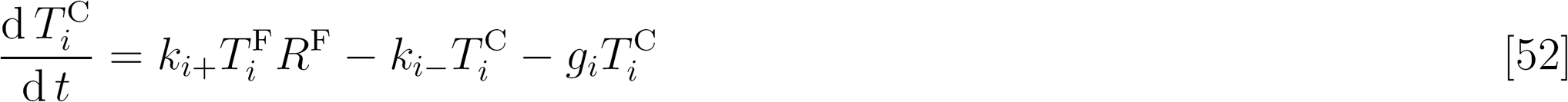

At steady states, by adding Eqs. 50 and 52, we get

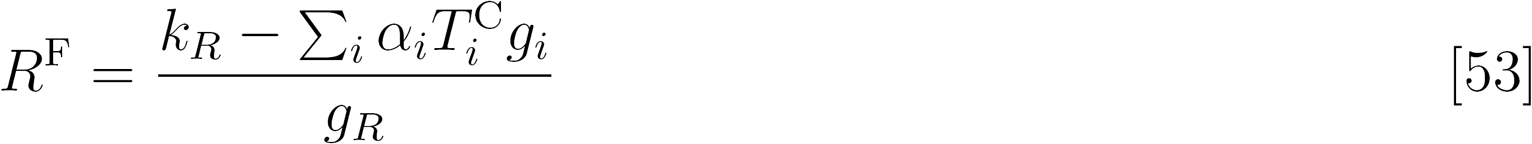

Solving Eq. 52, we get

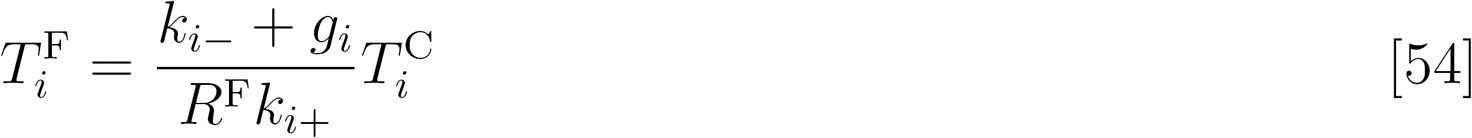

Combining Eqs. 53 and 54, we get

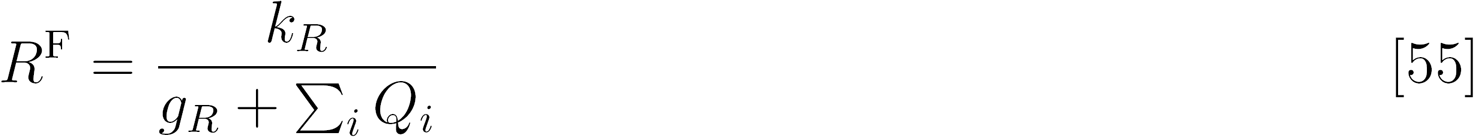

Where

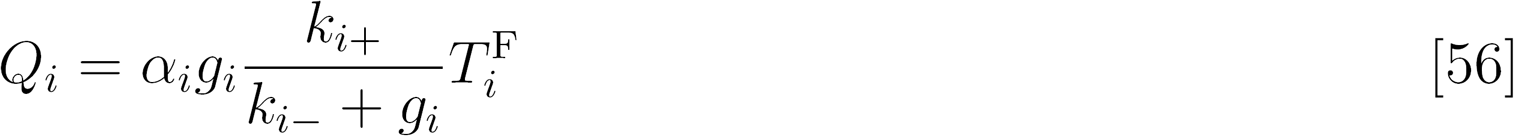

Thus,

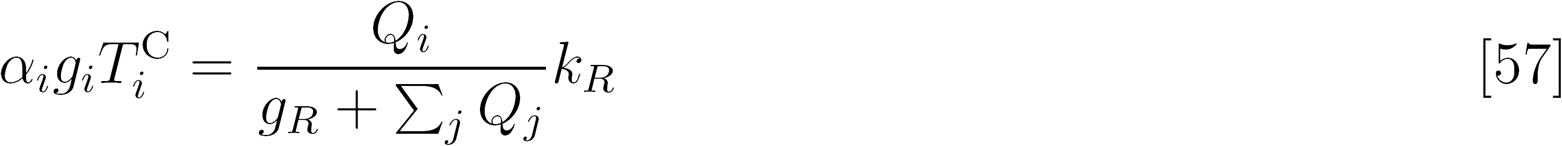

Eq. 57 has the exact form of current divider rule in electronics:

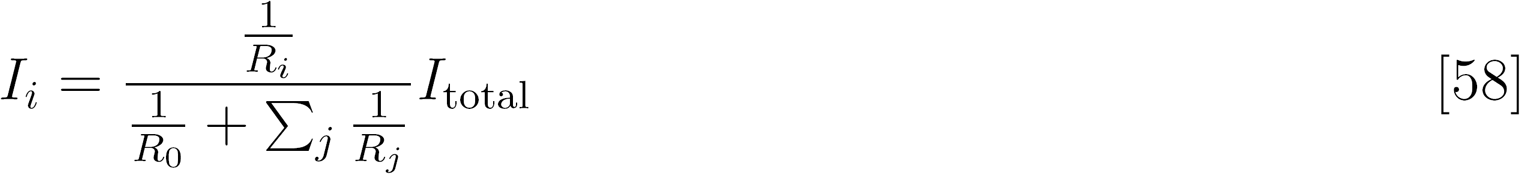

It inspires that *R*’s production rate (*k_R_*) is analogous to the total current (*I_total_*); the capability of 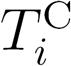 to consume 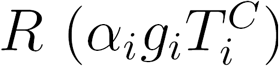 is analogous to the *i*th branch current (*I_i_*); and the capability of 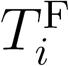 to occupy *R*(*Q_i_*) is analogous to the ith branch conductance (1/*R_i_*).

When *R* is scarce, 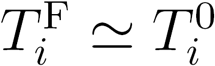, thus Eq. 56 is approximated to

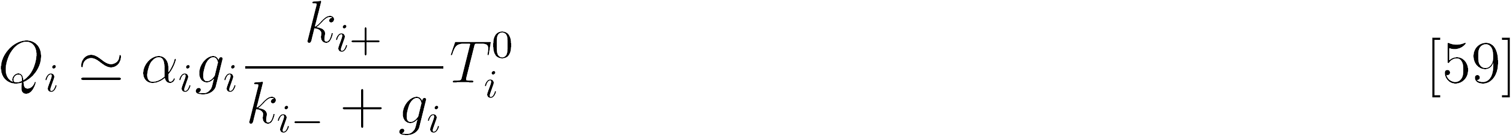

which indicates that in the “R scarce” regime, the capability of *T_i_* to occupy *R* (resistance) is only determined by the parameter settings of *T_i_*.

For catalytic reactions with a constat level of enzyme (regulator) and substances (targets), Eqs. 50-52 degenerate as

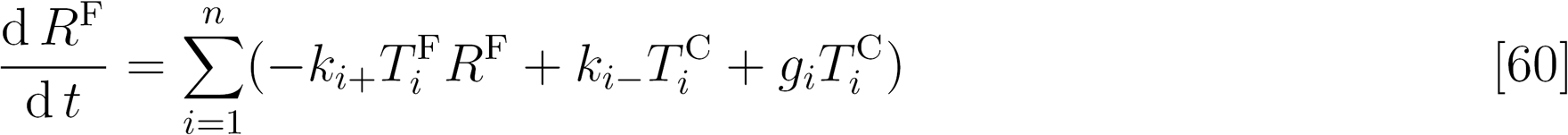

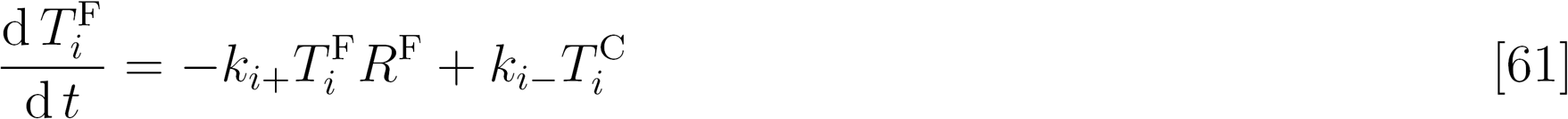

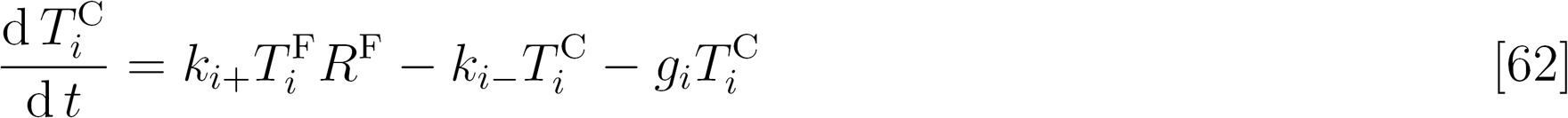

Under the assumption of Michaelis-Menten kinetics that 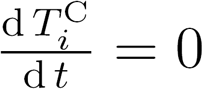 define *K_i_ = (k_i-_* + *g_i_)/k_i+_*, then we get

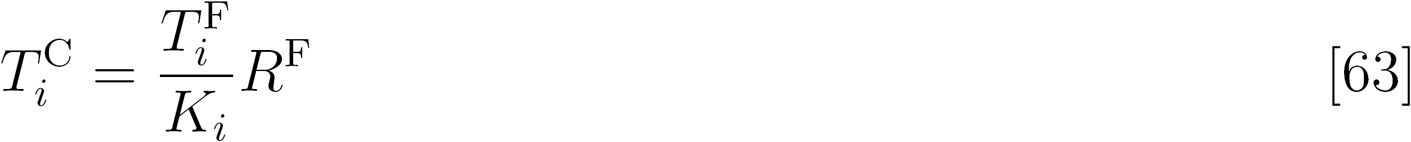

Thus,

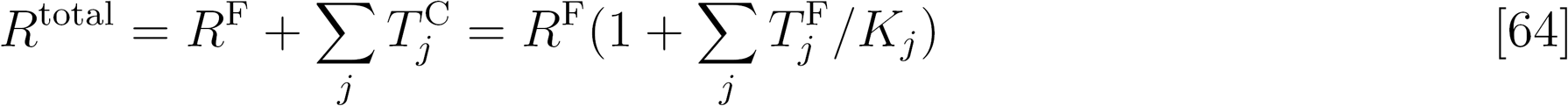

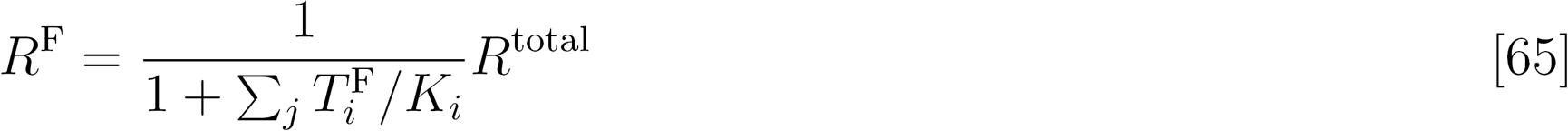

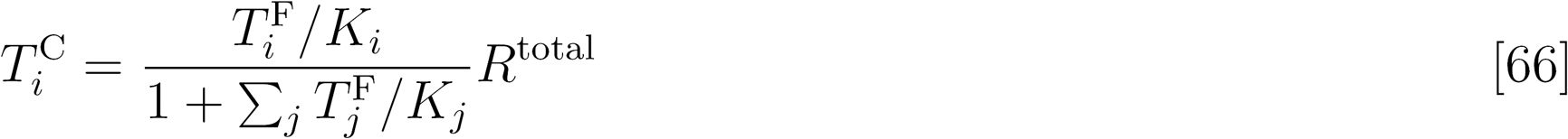

Which is the formation of enzyme allocation in Michaelis-Menten kinetics systems (Chou and Talaly, 1977).

### 7. Simulation parameters for drawing figures

The scales of simulation parameters are referenced from previous publications across different competition scenarios, such as transcription (Jayanthi et al., 2013), post-transcription (Ala et al., 2013; Schmiedel et al., 2015),translation (Gorochowski et al., 2016),degradation (Cookson et al., 2010) and chemical buffer solutions. Table S1 lists the parameters for drawing figures, the scales of which are derived from from previous researches on ceRNA effects (Ala et al., 2013; Yuan et al., 2015; Yuan et al., 2016)All gradually changing parameters are shown in figures. In Figure 2F-G and S2D-E, *k*_*T*2_ = 1 × 10^−2^. In Figure 2I, *k*_*T*2_ = 1 × 10^−4^, and parameters of *T*_3_ are shown in Table S1. In Figure 3A-C and S3A-F, *g*_1_ = 4 × 10^−5^, *k*_*T*2_ = 1 × 10^−4^. In Figure S3A, *g*_*T*2_ = 3.2 × 10^−4^. In Figure 3E-H, for dashed blue lines, *k*_*R*_ = 5 × 10^−3^; for thick blue lines, *k_R_* = 7.81 × 10^−5^; for thick green lines, *k_R_* = 5 × 10^−3^, *k*_*T*2_ = 3.21 × 10^−2^. In Figure S3G-J, *k*_*T*1_ = 5 × 10^−3^.

**Table S1.** Primary parameters for simulations.

| Primary parameters |  |  |  |  |  | Additional parameters<br>for Figure 2I |  |
| --- | --- | --- | --- | --- | --- | --- | --- |
| $R$ | | $T_1$ | | $T_2$ | | $T_3$ | |
| $k_R$ | $5 \times 10^{-3}$ | $k_{T1}$ | $1 \times 10^{-3}$ | $k_{T2}$ | $8 \times 10^{-3}$ | $k_{T3}$ | $5 \times 10^{-4}$ |
| $g_R$ | $1 \times 10^{-4}$ | $g_{T1}$ | $1 \times 10^{-5}$ | $g_{T2}$ | $1 \times 10^{-5}$ | $g_{T3}$ | $1 \times 10^{-5}$ |
| | | $k_{1+}$ | $1 \times 10^{-4}$ | $k_{2+}$ | $1 \times 10^{-4}$ | $k_{3+}$ | $1 \times 10^{-6}$ |
| | | $k_{1-}$ | $5 \times 10^{-5}$ | $k_{2-}$ | $5 \times 10^{-5}$ | $k_{3-}$ | $5 \times 10^{-5}$ |
| | | $g_1$ | $8 \times 10^{-5}$ | $g_2$ | $8 \times 10^{-5}$ | $g_3$ | $1 \times 10^{-5}$ |
| | | $\alpha_1$ | 1 | $\alpha_2$ | 0.5 | $\alpha_3$ | 0.5 |
| | | $\beta_1$ | 1 | $\beta_2$ | 1 | $\beta_3$ | 1 |
| | | $k_{P1}$ | $1 \times 10^{-2}$ | | | | |
| | | $g_{P1}$ | $5 \times 10^{-6}$ | | | | |

**Figure S1.**
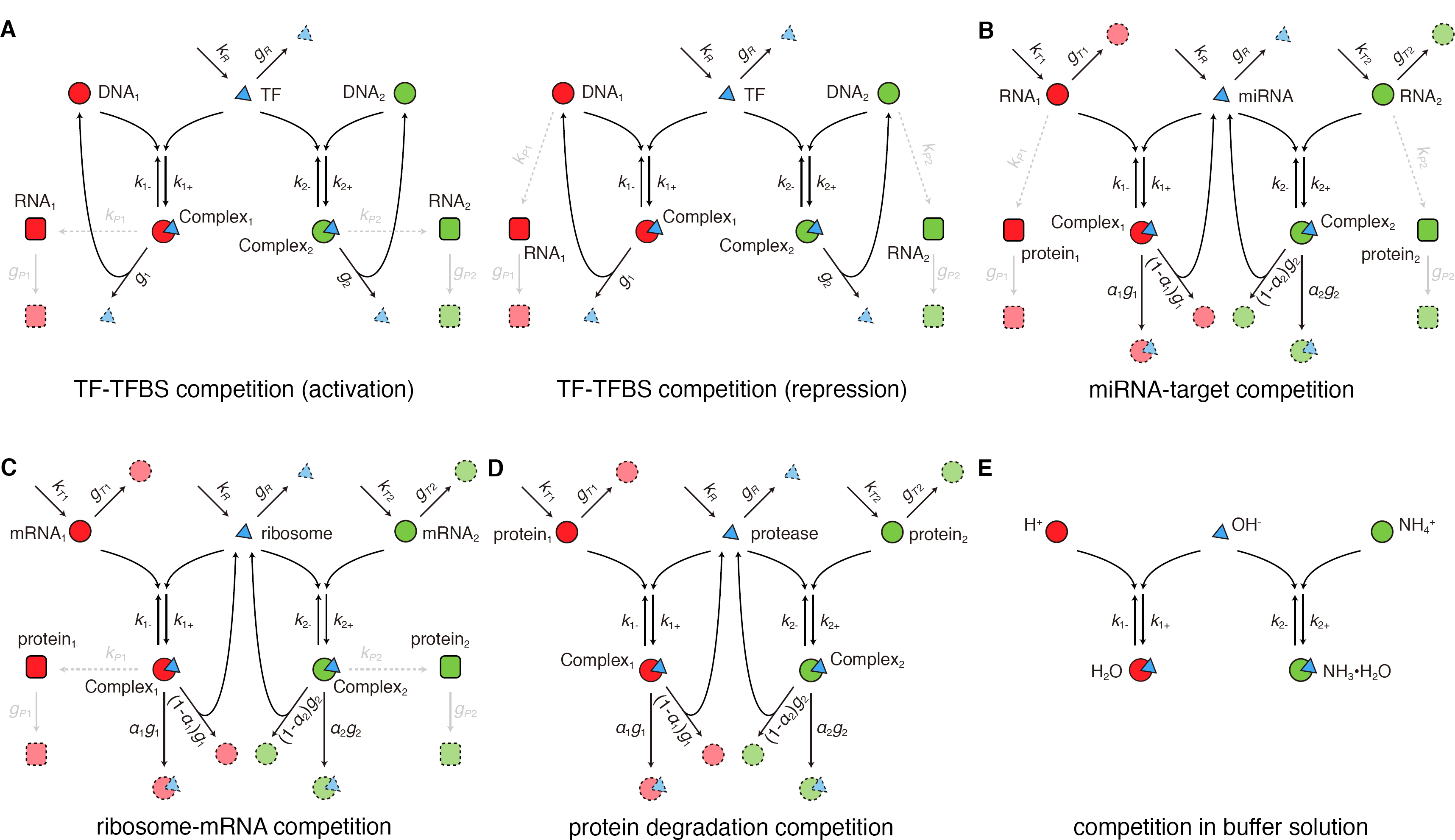
Detailed descriptions under the unified coarse-gained competition motif model for diverse competition scenarios. (**A**) DNA TF binding sites (TFBS) competing for TFs. Left: TF acts as an activator; right: TF acts as a repressor. (**B**) RNA molecules competing for miRNAs. (**C**) RNA molecules competing for ribosomes. (**D**) Proteins competing for proteases. (**E**) Competition in the ammonium buffer solution, where H^+^ and NH_4_^+^ compete for OH^-^.

**Figure S2.**
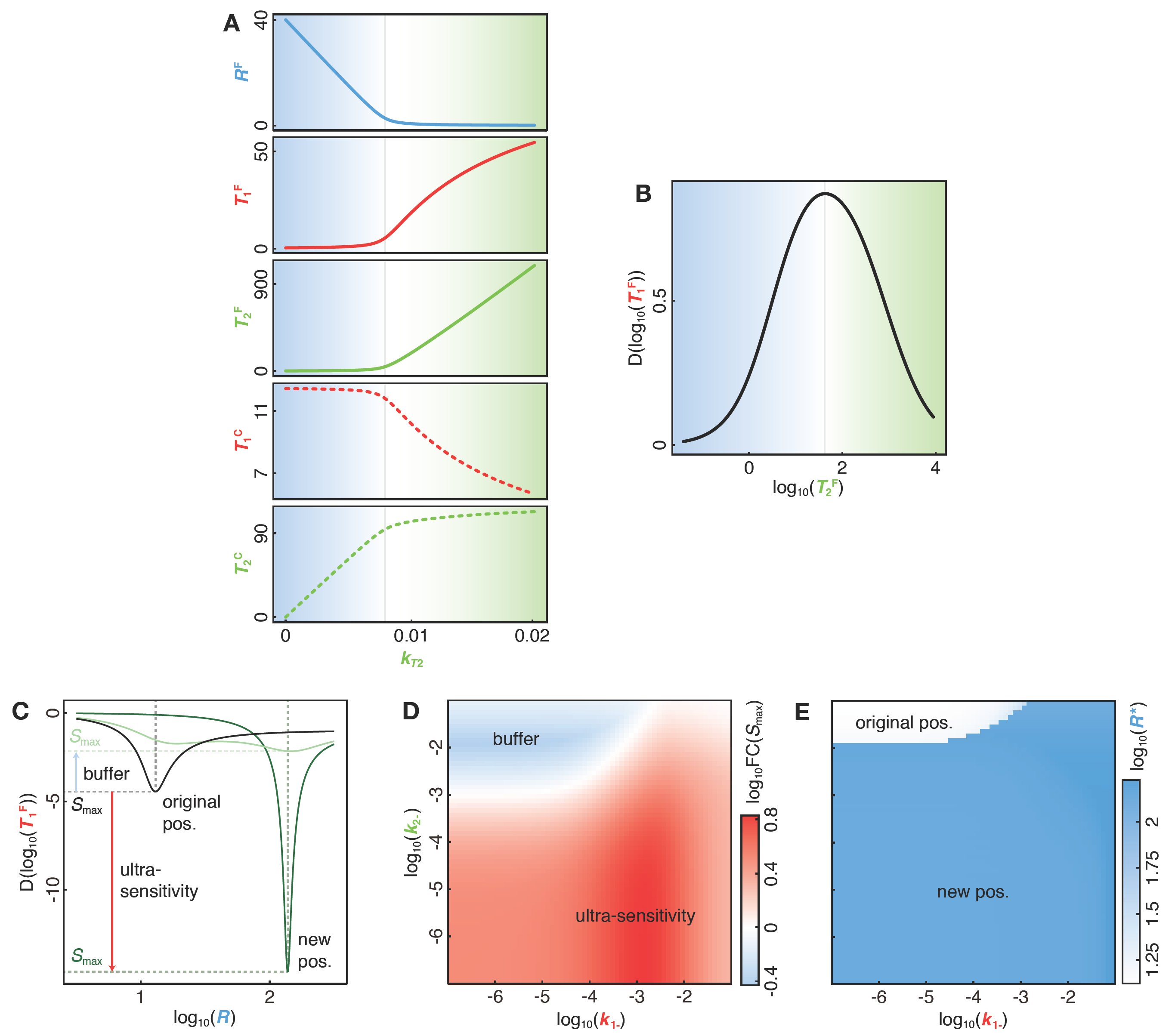
Steady state behaviors of competition systems. (**A**) Abundances of each component in Figure 2A on linear scales. (**B**) Derivatives of the curve in Figure 2B. Each component’s abundance in competition motif changing with *T*_2_’s production rate (*K*_*T*2_). Colors, lines and parameter settings are the same with Figure 2A-C. (**C**) Schematic diagram depicting the maximum sensitivity (S_max_) and its position (R*) of R-*T*_1_ dose-response curves. Dose-response curves are adopted from Figure 3G. (**D-E**) Relative binding affinities of *T*_1_ and *T*_2_ (*k*_1_-and *k*_2_-) shape *R*-*T*_1_ dose-response curves. (**D**) Fold change of Smax compared with that of non-competing system (*k*_*T*2_=0). Competition of *T*_2_ buffers the response of *T*_1_^F^ to *R* when log_10_FC(S_max_)<0, but introduces larger sensitivity when log_10_FC(S_max_)>0. (**E**) Changes of R*.

**Figure S3.**
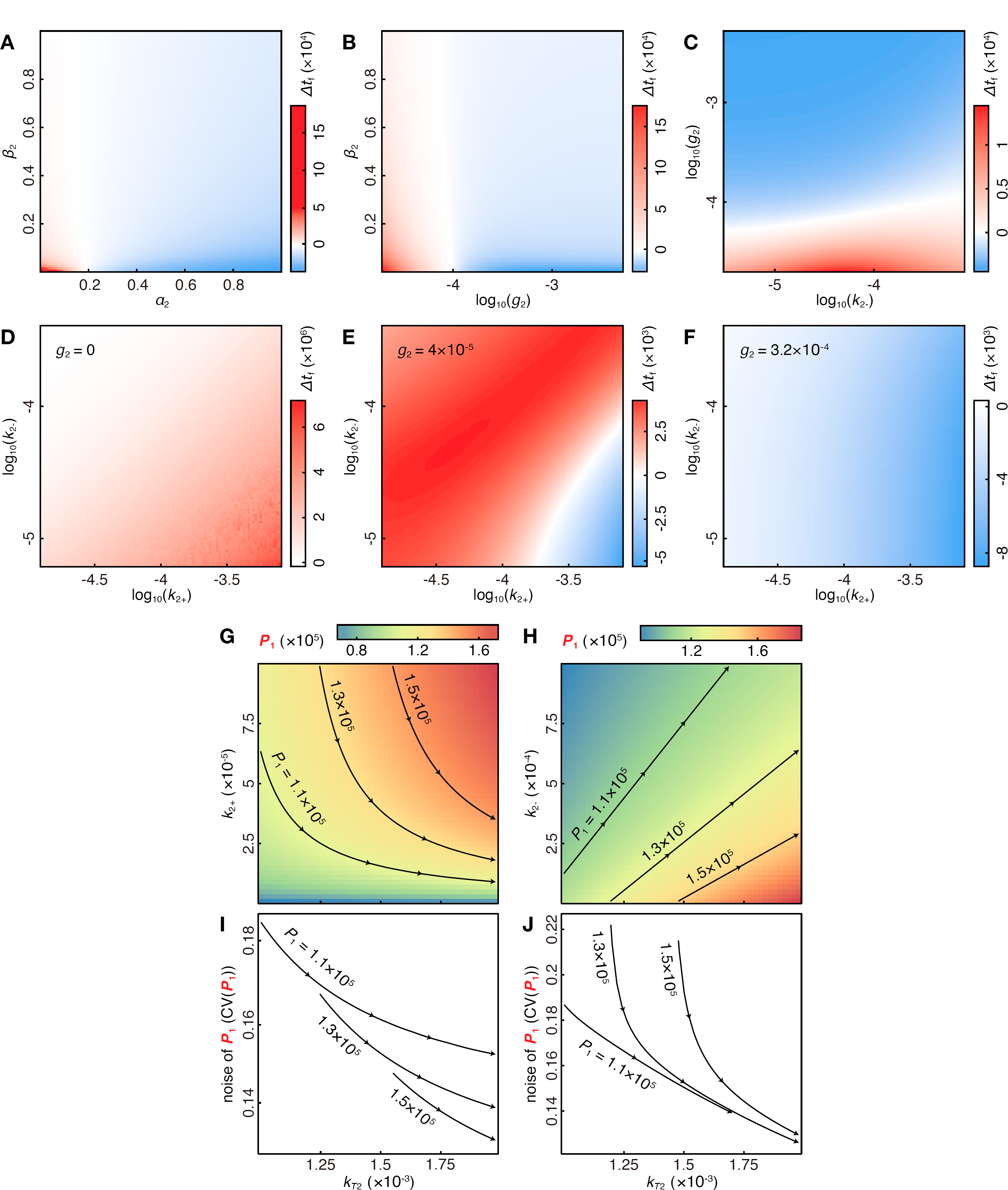
Dynamic properties of competition systems. (**A-F**) Modifications of response time on the falling edge of *R*’s change under different kinetic parameters: (**A**) different *α*_1_ and *β*_2_; (**B**) different *g*_2_ and *β*_2_; (**C**) different *k*_2-_ and *g*_2_; (**D-F**) different *k*_2+_, *k*_2-_ and *g*_2_ (values of *g*_2_ are shown in figures). (**G-J**) Abundant weak competitors can buffer target expression noise better. (**G-H**) *P*_1_ level changing with *T*_2_’s production rate (*k*_*T*2_) and *T*_2_’s association rate (*k*_2+_) (**G**), or *k*_*T*2_ and *T*_2_’s dissociation rate (*k*_2-_) (**H**). Black lines are *T*_1_^F^ level isolines. (**I-J**) Target expression noise changes on the isolines in (**G**) and (**H**) respectively. Along the direction of the arrows, *T*_2_’s production increases and *T*_2_’s binding affinity decreases, bringing about lower expression noise.

